# Direct and indirect striatal projecting neurons exert strategy-dependent effects on decision-making

**DOI:** 10.1101/2024.04.15.589515

**Authors:** Elena Chaves Rodriguez, Jérémie Naudé, Daniel Rial, Alban de Kerchove d’Exaerde

## Abstract

The striatum plays a key role in decision-making, with its effects varying with anatomical location and direct and indirect pathway striatal projecting neuron (d- and i-SPN) populations. Using a mouse gambling task with a reinforcement-learning model, we described of individual decision-making profiles as a combination of three archetypal strategies: Optimizers, Risk-averse, Explorers. Such strategies reflected stable differences in the parameters generating decisions (sensitivity to the reward magnitude, to risk or to punishment) derived from a reinforcement-learning model of animal choice. Chemogenetic manipulation showed that dorsomedial striatum (DMS) neurons substantially impact decision-making, while the nucleus accumbens (NAc) and dorsolateral striatum neurons (DLS) have lesser or no effects, respectively. Specifically, DMS dSPNs decrease risk aversion by increasing the perceived value of risky choices, while DMS iSPNs emphasize large gains, affecting decisions depending on decision-making profiles. Hence, we propose that striatal populations from different subregions influence distinct decision-making parameters, leading to profile-dependent choices.

## INTRODUCTION

To adapt to dynamic environments, decision processes aim to select the course of action that will lead to the best outcome. Value-based decision-making can be described as the iterative process of selecting actions on the basis of their expected values and of evaluating outcomes to update action values(*1*, *2*). Decision-making is therefore fundamentally based on predictions of the probability of obtaining the different available outcomes (e.g., rewards, omissions or punishments), as well as estimations of their respective values. There has been a renewed interest in understanding how the nervous system weights the expected values of outcomes to choose between options.

Decision-making processes emerge from the mesocorticolimbic loop, which has a key, yet not fully understood, role in the striatum (*3–5*). The rodent striatum is divided into several regions, each of which is thought to have a particular function in decision-making. The dorsal striatum is responsible for motor and cognitive control associated with goal-directed actions (DMS) or the generation of habits (DLS), and the NAc (*6–15*) manages reward, motivation and Pavlovian associations. While habits, which consist of stimulus‒response associations, are generally insensitive to the value of the outcome (*16*, *17*), Pavlovian (stimulus‒outcome) and goal‒ directed (action‒outcome) associations can affect value‒based decisions (*13*, *16*). How these associations with outcomes (rewarding or punitive) rely on striatal computations has been the subject of much study (*18*). Striatal projection neurons (SPNs) account for 95% of all striatal neurons. There are two distinct SPN subpopulations: dopamine D1 receptor (D_1_R)-expressing or direct SPNs (dSPNs) and dopamine D2 receptor/adenosine A2A receptor (A_2A_R)-expressing indirect SPNs (iSPNs). It remains debated how these two neuronal groups interact during decision-making (*19*). Loss-of-function or gain-of-function experiments (*20–25*) support a go/no-go model (*26*, *27*) in which dSPNs play a prokinetic role and iSPNs play an antikinetic role. However, correlative approaches (*28–33*) revealed the coactivation of dSPNs and iSPNs during motor performance. This suggests a complementary encoding of behaviors, with congruent activations observed in dSPNs for multiple behaviors and iSPNs for suppressing competing behaviors (*34*).

In the context of decision-making, dSPNs and iSPNs may encode potential rewards and costs (*19*, *35*), respectively, or encode and update values in goal-directed learning (*36*). However, manipulations of striatal dopamine also affect risk seeking (*9*), which does not fit into a clear reward/cost dichotomy, and the relative influences of dSPNs and iSPNs on risk seeking remain to be assessed. Moreover, the attitude toward risk, as well as the sensitivity to rewards, costs, or punishments, is highly idiosyncratic (i.e., it reflects stable individual traits). Different decision profiles have been described in humans and animals. Interindividual variability in the choice behavior of healthy individuals has been described in humans, rats (*37–39*) and mice (*40–42*). How such differences are related to variations in mesocorticostriatal properties remains an open question. Furthermore, several substances, ranging from caffeine to cocaine, can have different effects on individuals according to their choice behavior. For example, impulsive choice has been recurrently described as a predictor of susceptibility to developing addictive disorders in humans, such as alcohol or cocaine use disorders (*43–46*). Risk-preferring rats are more prone to develop cocaine addiction and are also more sensitive to cocaine craving during withdrawal (*47*). Additionally, in rats, caffeine, an agonist of A_2A_R, can have a motivational effect on performing effortful tasks in low performers, whereas in naturally motivated rats, caffeine can disrupt effortful performance (*48*). These pathological cases highlight that the effect of modifying the weight of a given decision variable (e.g., reward size) depends on the sensitivity to the other decision variables (e.g., risk attitude, sensitivity to effort). In this framework, the different roles of dSPNs and iSPNs from subparts of the striatum might thus depend upon the decision-making traits of the animals.

Here, we used a rodent Iowa Gambling Task (IGT) adapted from Young et al. (*49*) to show how different decision-making profiles arise in mice. Furthermore, we targeted dSPNs and iSPNs in the DMS, DLS and NAc with a chemogenetic tool to assess how decision-making strategies can be differentially altered by changes in striatal excitability, depending on “basal” decision-making traits. Computational analyses allowed us to identify three different decision-making archetypes in mice (Explorers, Risk-averse and Optimizers). These cognitive profiles were characterized by distinct sensitivities to risks, the subjective utility of large rewards, and explore-exploit trade-offs. We then showed that increasing DMS excitability exerted the most profound effects on choices and motivation, which is consistent with the IGT being a goal-directed task. Specifically, we found that facilitating DMS dSPN activity decreased safe choices in all the mice by decreasing risk aversion, whereas facilitating DMS iSPN activity decreased safe choices only in the Optimizer mice through a decrease in reward saturation. Compared with their DMS counterparts, increasing NAc dSPN excitability induced similar, albeit blunted, effects. NAc iSPN and DLS i- and d-SPN manipulations had nonspecific effects on motivation but not on choice behavior. Overall, we highlighted how the striatal subpopulation exerts cognitive profile-dependent effects on choice behavior.

## RESULTS

### Mouse gambling strategies arise from varying sensitivities to task parameters

To assess decision-making in a complex environment, we used a rodent-adapted version of the IGT (*49*), which we adapted for mice. The task takes place in an operant chamber in which animals can select among four nose-poke holes to obtain food pellets in a magazine on the opposite wall (Figure 1A, see Methods). As mice need to nose-poke on the holes between 5 s (shorter responses were not rewarded and counted as premature) and 10 s (longer responses were not rewarded and counted as omissions), this test measures both motivation/impulsivity and choice behavior. Each hole (P1-P4) is associated with a distinct reward probability *p* to obtain a reward magnitude (number of pellets) and a related probability of 1-*p* to obtain a punishment in the form of a time-out (TO). TOs correspond to a potential loss of reward because of the overall limit of time of the task, e.g., potential loss = (time-out duration × average pellet rate) × time-out probability, with the pellet rate taken as the number of pellets for the option divided by the task duration. As the potential (expected) gain is the number of pellets × reward probability, one can compute the (linear) “expected net return” for each option as follows: *E*(*X*_*i*_) = *p*_*i*_*R*_*i*_ − (1 − *p*_*i*_)*T*_*i*_ where *p*_*i*_ is the probability of reward, *R*_*i*_the reward magnitude, and *T*_*i*_ the punishment due to time-out, in terms of opportunity cost of time (time-out duration times the average number of pellet/s). This gives E(P1) = 0.89 pellets, E(P2) = 1.52 pellets, E(P3) = 0.60 pellets, E(P4) = −0.32 pellets. In this context, options P1 and P2 are labeled as “safe choices” because they deliver a higher net return through a smaller amount of reward than P3 and P4 but with a higher probability and shorter associated TOs. In the original context of the task(*40*, *50*, *51*), P3 and P4 are thus considered maladaptive: despite delivering more pellets on rewarded trials, the high probability of a long time-out renders them disadvantageous in the long run (see Methods). In our experiments, the mice indeed preferred, on average, the safe options (P1 and P2 compared with P3 and P4), and such choice behavior was stable across sessions (Figure 1B; repeated-measures (RM-)ANOVA: time: F_(3,668)_=92.2, p<1e^−12^; options F_(3,668)_=0.1, n.s.). However, such average preferences often mask important variability. Hence, we leveraged the analysis of baseline (before chemogenetic/ Designer Receptor Exclusively Activated by Designer Drugs (DREADD) manipulations, see below) choice behavior from all the mice (n=168) to characterize such variability. There was marked variability both in terms of choice behavior, with P1 (the safest option) chosen as little as 0% or as much as 100% of the time by a given animal (Figure 1C), and in terms of motivation/impulsivity (number of trials, % of premature responses, and % of omissions; Figure 1D). We thus sought to delve into the origin of such variability.

**Figure 1.**
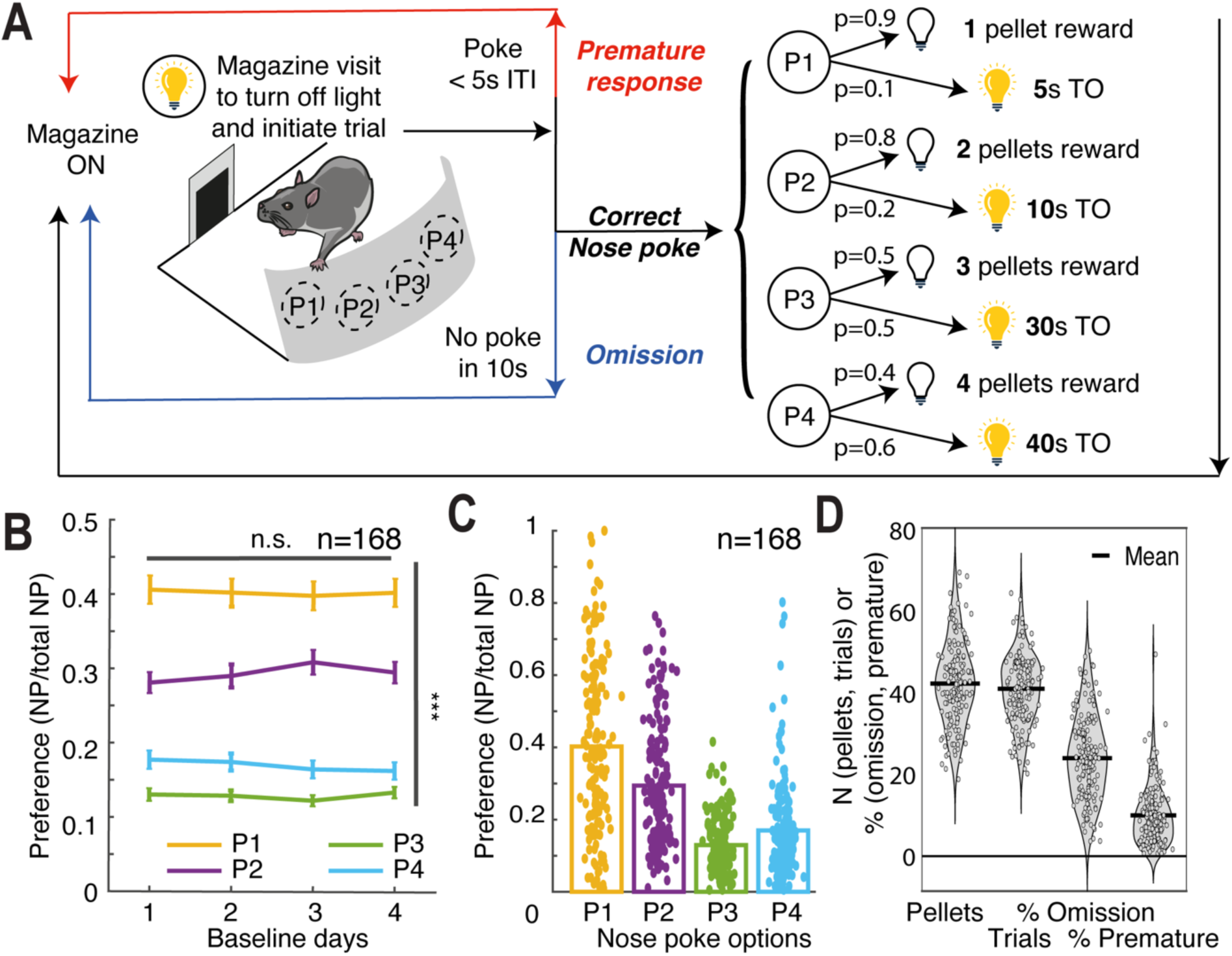
Mice exhibited different but stable preferences in the gambling task. **A.** Task schema: upon initiating a trial by magazine entry, the mice had to choose. **B.** Preferences for the four options against days of training, showing that the mice learned the task and displayed a stable preference for the safe options. **C.** The preferences of all mice (all conditions) displayed substantial interindividual variability. **D.** The other behavioral measures in the gambling task (number of trials, proportions of premature responses, and omissions) also displayed substantial interindividual variability, suggesting different choice strategies. n.s. not significant; *** p<0.001.

A classical way to describe interindividual variability is by introducing a dichotomy between “good” (or “safe”) and “poor” (or “risky”) decision-makers. We used such a distinction (mice were considered “safe” if they had a preference for the advantageous options P1 and P2) so that our characterization of the roles of striatal subpopulations could be compared with other studies using a rodent gambling task. However, such categorizations hinge on the experimenter’s predefined criteria for what is considered beneficial or detrimental to the animals. Moreover, dichotomies resulting from arbitrary thresholds to partition data may overlook the nuanced statistical intricacies of interindividual variability. Consequently, in addition to the safe/risky dichotomy, our approach aimed to identify behavioral profiles that not only captured (1) the statistical patterns within the behavioral data but also reflected (2) the generative properties of the underlying decision-making architecture.

As a starting point, we observed strong correlations between the different individual measures (choice behavior and motivation/impulsivity), e.g., the tendency to choose the safe choices (P1 and P2) was negatively correlated with the proportion of omissions (R^2^ = 0.27, p=6.10^−13^, Figure 2A, Supp. Figure 1A-C). This statistical structure hints at constraints in the mechanisms generating the data (i.e., the decision-making traits generating the animal behavior). We thus reduced the high-dimensional dataset (7 measures: preferences for the 4 options, i.e., P1-P2-P3-P4, as well as the number of trials, omissions and premature responses) via principal component analysis (PCA; Supp. Figure 1D). As there were no clearly separated clusters, we expressed individuals as a continuum between “archetypal” strategies via archetypal analysis, as previously developed (*52*, *53*) and applied to decision-making (*41*). Compared with classical clustering, which groups individual data around typical observations (cluster centers) that constitute a decision-making profile, archetypal analysis depicts individual behavior as intermediates between extreme strategies, i.e., archetypes (Figure 2B). This framework acknowledges that, contrary to experimenter-defined performance in each task, animals define their own success criteria, generally as a tradeoff between different objectives. Indeed, no decision-making profile (being averse to risk, optimizing the expected reward, showing a low level of attentional control) can be optimal for all the possible environments (in terms of reward scarcity, risk of punishment, etc.) the animals might face. The best compromises between objectives lead to phenotypes that lie in low-dimensional polytopes in the trait space (*52*, *53*). Archetypal strategies corresponding to the objectives lie at the apices (extrema) of these polytopes, and individual strategies are described as linear combinations of the archetypes. If the behavior in the mouse gambling task indeed emerges as an individual solution to a trade-off between archetypal strategies, then such a description should 1) be stable across sessions and 2) reflect underlying decision-making traits.

**Figure 2.**
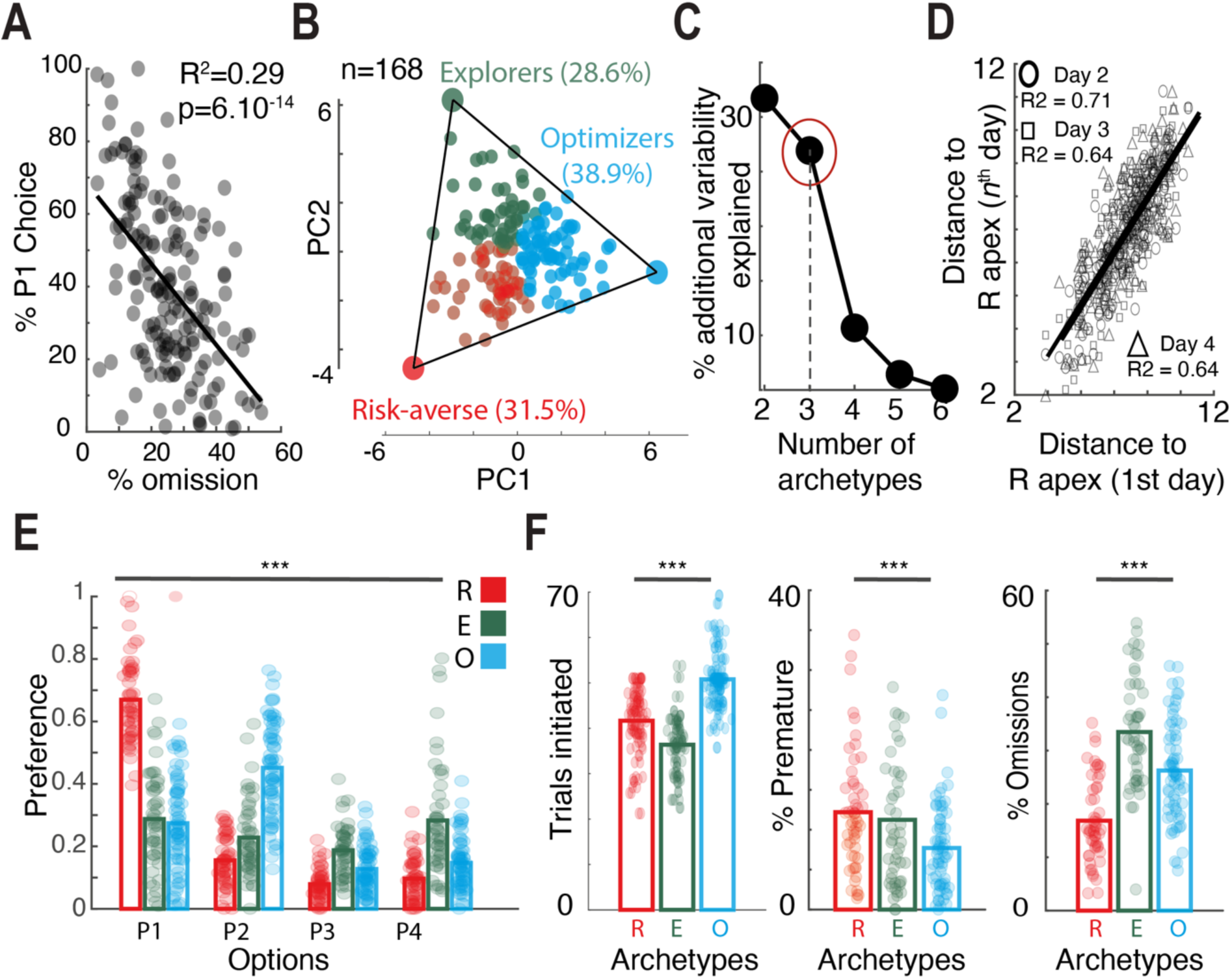
Variability in mouse behavior is a continuum between archetypal strategies. **A.** Preference for the safe choice was correlated with the percentage of omissions. **B**. Archetypal analysis performed on 8 measures (number of trials, pellets obtained, % omission, % premature responses, and preferences for the 4 options) with n = 168 mice. Each point of the ternary plot represents the projection of an individual onto the two principal components (PCs) derived from the 8 measures. The apices of the polytope encompassing the data define the archetypal strategies (here, a triangle with 3 archetypes: Optimizers (O), Risk-averse (R), Explorers (E)). Individuals can be described as convex combinations of the 3 archetypes. **C.** Variability (in the 8 measures × 168 mice) explained as a function of the number of archetypes, showing that 3 archetypes constitute a parsimonious description of the data. **D.** The distance to archetypes (R apex) was highly correlated between consecutive days, suggesting that behavior in the task arose from stable cognitive profiles. **E**. Preferences from mice assigned to their closest archetype indicate that P1 (certain, small reward, short TO) was preferred by Risk-averse (R) mice, and that Optimizers (O) preferred P2 (larger, less certain reward, more advantageous relative to TO than P1), whereas Explorers (E) did not exhibit a preference. **F**. These choice strategies were associated with differences in other task measures: Risk-averse mice displayed fewer omissions, Optimizers earned more pellets and made fewer premature responses, and Explorers made more omissions and earned fewer pellets. *** p<0.001.

We first calculated the number *n* (n being less than the dimension of the data) of archetypes by fitting polytopes with *n* apices to the data and chose *n=3,* beyond which there was little improvement in the explained variance (Figure 2C, Supp. Figure 1E, F). The archetypes positions in the PCA space were robust to the use of only a subset of the data (jackknife resampling, Supp. Figure 1G). The archetype analysis explained more variance than did the classical safe/not safe dichotomy (Supp. Figure 1H, I). We then verified that individual strategies were stable by computing the distance of individual animals (expressed as coordinates in the PCA space) to the apices (the archetypes). This distance-to-archetype was highly correlated between consecutive days (Day 1 versus Day 2: R^2^= 0.71, p=3.10^−46^; Day 1 versus Day 3: R^2^= 0.64, p=3.10^−38^; Day 1 versus Day 4: R^2^= 0.64, p=8.10^−39^, n = 169 mice, Figure 2D, Supp. Figure 1J, K), indicating stable profiles. Finally, partitioning the animals into three groups, on the basis of their proximity to one of the archetypes, corresponded to significant differences in the behavioral data (choices: 2-way ANOVA: F_(6,66)_=115.86, p=1.10^− 99^ ; trials: 1-way ANOVA, F_(2,165)_=69.82, p=1.10^−22^; premature responses: 1-way ANOVA, F_(2,165)_=41.68, p=1.10^−15^; omissions: 1-way ANOVA, F_(2,165)_=5.92, p=3.10^−3^). The first group (hereafter, “Risk-averse”, R) displayed a greater preference for P1, i.e., the least risky option (Figure 2E), more premature responses, and fewer omissions (Figure 2F), than did the other two groups. The second group (“Explorers”, E mice) displayed less marked preferences (the flatter pattern of choices in Figure 2E), initiated fewer trials, and made more omissions (Figure 2F). The third and last group (“Optimizers”, O mice) preferred P2, i.e., the more advantageous option, over P1 (Figure 2E), initiated more trials, and made less premature (impulsive) responses (Figure 2F).

The fact that behavioral features were enriched at each archetype is only confirmatory (like significant differences between subgroups following a median split) because an archetypal analysis is performed on the PCA space derived from the same behavioral data. We thus sought to confirm that the statistical distribution in the “manifested” choices of the animals arises from differences in underlying (“hidden”) decision-making strategies. To test for putative differences in their subjective valuation/decision processes, we fit the animals’ choices with a reinforcement-learning model (Figure 3A). In this model, the probability of choosing the hole Px (e.g., P1) depends on its expected value compared with the sum of values from all the other options. The model assumes that the value of an option depends on its expected (subjective) reward, punishment, and risk (see the Methods for details). The expected reward (or punishment) equals the reward (or punishment) probability times the magnitude of the subjective reward (or punishment), such as for the “expected net return”. Nevertheless, a major difference between the IGT in rodents and those in humans is that the reward takes the form of food pellets (instead of points or money), and punishments are time-outs. Hence, classical models considering that animals compute an “expected net return” rely on the hypothesis of linear preferences (or time perception), e.g., that winning 4 pellets (or waiting 40 s) from one choice is similar to having 4 times 1 pellet from 4 choices (or waiting 4 times 10 s, respectively). In contrast, animals often display nonlinear subjective values or time perceptions. Hence, we introduced an additional parameter ρ that depicts how the subjective value saturates with the number of pellets: ρ<1 corresponds to a value of 4 pellets lower than 4 times the value of 1 pellet, henceforth favoring small gains (i.e., a small ρ favors small gains, whereas a high ρ favors large gains). Similarly, the parameter *T* depicts how subjective punishment depends on the time-out duration. The option value also incorporates a risk-sensitive term (φ) that can be positive (risk prone) or negative (risk averse). Finally, we included an inverse temperature parameter (β) to account for the exploitation–exploration tradeoff: animals choosing their preferred option nearly all the time were considered exploitive (high β), whereas animals with a “flat” distribution of choices (no marked preference) were labeled explorative (low β). We checked that this model provided a better fit than simpler models used in the literature (Figure 3B); in particular, nonlinear reward and time perception resulted in a better fit (Figure 3B, Supp. Figure 2A) and generation (Figure 3C) of choice patterns. We also verified that model parameters could be recovered with the relatively small number of trials performed by the animals, demonstrating a reasonable level of accuracy (Supp. Figure 2B). By fitting this model to individual data, we could express the choices from each animal as if it had been generated by a decision-maker with a given set of parameters (β, ρ, *T*, φ; Figure 3C). This decision model allowed us to better characterize the archetypal strategies (Figure 3D, β parameter: 1-way ANOVA, F_(2,165)_=13.67, p=3.10^−6^; ρ parameter: 1-way ANOVA, F_(2,165)_=15.73, p=6.10^−7^; *T* parameter: 1-way ANOVA, F_(2,165)_=8.83, p=2.10^−4^; φ parameter: 1-way ANOVA, F_(2,165)_=24.49, p=4.10^−10^). The R group displayed the lowest φ parameter (indicating strong aversion to risk) but also a high sensitivity to time-out *T* and a low sensitivity to reward ρ. The E group was characterized by a low inverse temperature β, indicative of high exploration (or low exploitation), in line with its absence of a clear preference among the four options. The O group had more balanced decision parameters, with a high β (exploitive), an insensitivity to risk on average, and near linear saturation functions (ρ and *T* close to 1) for reward and punishment. Expressing individual fits as intermediates between the archetypes extrema yielded the same interpretation (Supp. Fig 3A). suggesting that the computational characterization does not arise from partitioning the data into clusters. The conjunction of the archetypal analysis with the computational model further suggests that the stable interindividual variability observed in the mouse gambling task reflects stable (Supp. Figure 3B) decision-making profiles.

**Figure 3.**
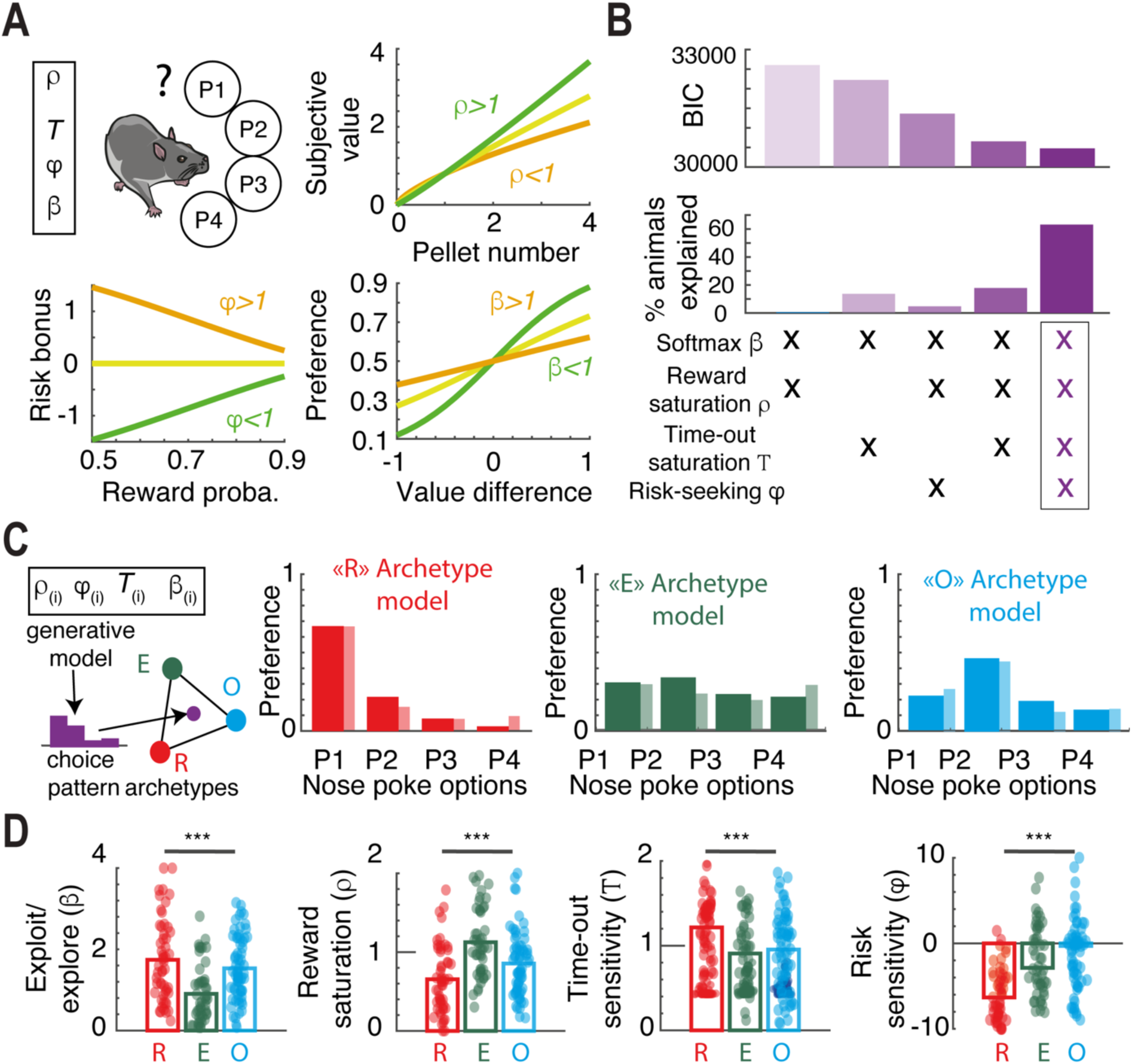
Archetypal behaviors arise from different decision-making strategies. **A**. Sketch of the computational model of decisions (“choice model”): Mice were assumed to assign a value to each of the four options (P1-P4) depending ons how much they positively valued a pellet (ρ parameter), how much they considered the time-outs to be punishing (*T* parameter), their attitude toward risk (i.e., toward outcome probabilities closer to 0.5, φ parameter) and the precision in their decision process (“inverse temperature” β parameter). **B**. Bayesian Information Criterion (top, model likelihood penalized for model complexity) and proportion of animals best explained (middle) for different versions of the Reinforcement-Learning model (bottom), with the 4-parameters model showing smaller (i.e. better) BIC. **C**. Average preferences for the 4 options (P1-P4) for the different archetypes (O for Optimizers, R for Risk-averse, E for explorers) can be generated through different combinations of decision-making parameters (β, ρ, *T*, and φ). Opaque bars display the model fit and shaded bars display the experimental data. **D.** The choice model suggests that the strategies unraveled by the archetypal analysis arise from different sensitivities to task variables: mice were in the Risk-averse archetype if their sensitivity to punishment was higher and their attitude toward risk was more negative; Explorers mainly displayed lower β values (low precision, or high noise, in the decision process); and Optimizers exhibited higher sensitivity to reward and their attitude toward risk was neutral. *** p<0.001.

### Dorsomedial dSPNs decrease safe choices by promoting risk-seeking

We next sought to assess the contributions of both SPN populations to decision-making in the main striatal subdomains (DMS, DLS and NAc). We thus used DREADD activation, rather than optogenetic activation (see Discussion section), to facilitate neuronal activity from a specific subpopulation after the values of the different options were learned by the animal to focus on the decision process. For this purpose, we tested 6 different groups of mice, one per SPN population (direct or indirect pathway SPNs) and striatal area (DMS, DLS or NAc). We bilaterally injected a Cre-dependent hM3Dq DREADD virus (AAV5 hSyn-DIO-hM3D(Gq)-mCherry) or a control fluorophore lacking the hM3Dq construct (AAV5 hSyn-DIO-mCherry)s into the striatal zone of interest (DMS, DLS or NAc) in D_1_R-Cre (*54*) or A_2A_R-Cre mice (*22*) to selectively express hM3Dq or mCherry on either dSPNs or iSPNs, respectively (Supp. Figure 4).

As the mouse gambling task relies on instrumental behavior, we first focused on DMS d-SPNs (Figure 4A). Given that mice received DREADD ligand (CNO, 1 mg/kg) injection on the last day of behavioral testing (see below), we verified ex vivo the effectiveness of the hM3Dq DREADDs in increasing the intrinsic excitability of targeted neurons by collecting slices from mice (hM3Dq and mCherry controls) just after the behavioral testing. The marker of neuronal activation Fos was colocalized with mCherry expression only in hM3Dq animals (Figure 4B, Supp.Table 1, t_(16)_=8.5, p<0,0001; unpaired t test), confirming that CNO facilitated neuronal activation during the mouse gambling test. Next, we measured that neuronal activation was followed by an increase in intrinsic excitability (i.e., active membrane properties), as expected from hM3Dq recruitment (*55*). We did not find any changes in the passive properties of the neurons via patch-clamp recordings: the resting membrane potential did not change following CNO injection (Supp. Figure 5A, t_(10)_=0.2, n.s.). We did not observe any modifications in (putatively corticostriatal or thalamicostriatal) synapse strength as measured by the AMPA/NMDA ratio (t_(10)_=0.93, n.s., Supp. Figure 5B), suggesting that Fos activity was not due to broad changes in synaptic excitation. Owing to the intrinsic excitability of the cells, the rheobase (minimal current needed to elicit an action potential) was not changed following hM3Dq + CNO treatment (t_(10)_=0.96, n.s., Figure 4C left). However, the number of action potentials elicited by intermediate current intensities increased following hM3Dq + CNO treatment, indicating an increase in neuronal gain (in the linear range of the frequency intensity, or f-I curve, 2-way ANOVA F_(14,140)_ = 13, p<0.0001, Figure 4C right). This finding suggests that neuronal Fos activation arose in neurons receiving a notable drive (putatively task-related) amplified by increased neuronal gain rather than from nonspecific electrical activity (which would follow from changes in resistance or rheobase). Together, the patch clamp recordings confirmed that CNO injection before behavioral testing had lasting effects (detectable ex vivo after the session) on neuronal excitability.

**Figure 4.**
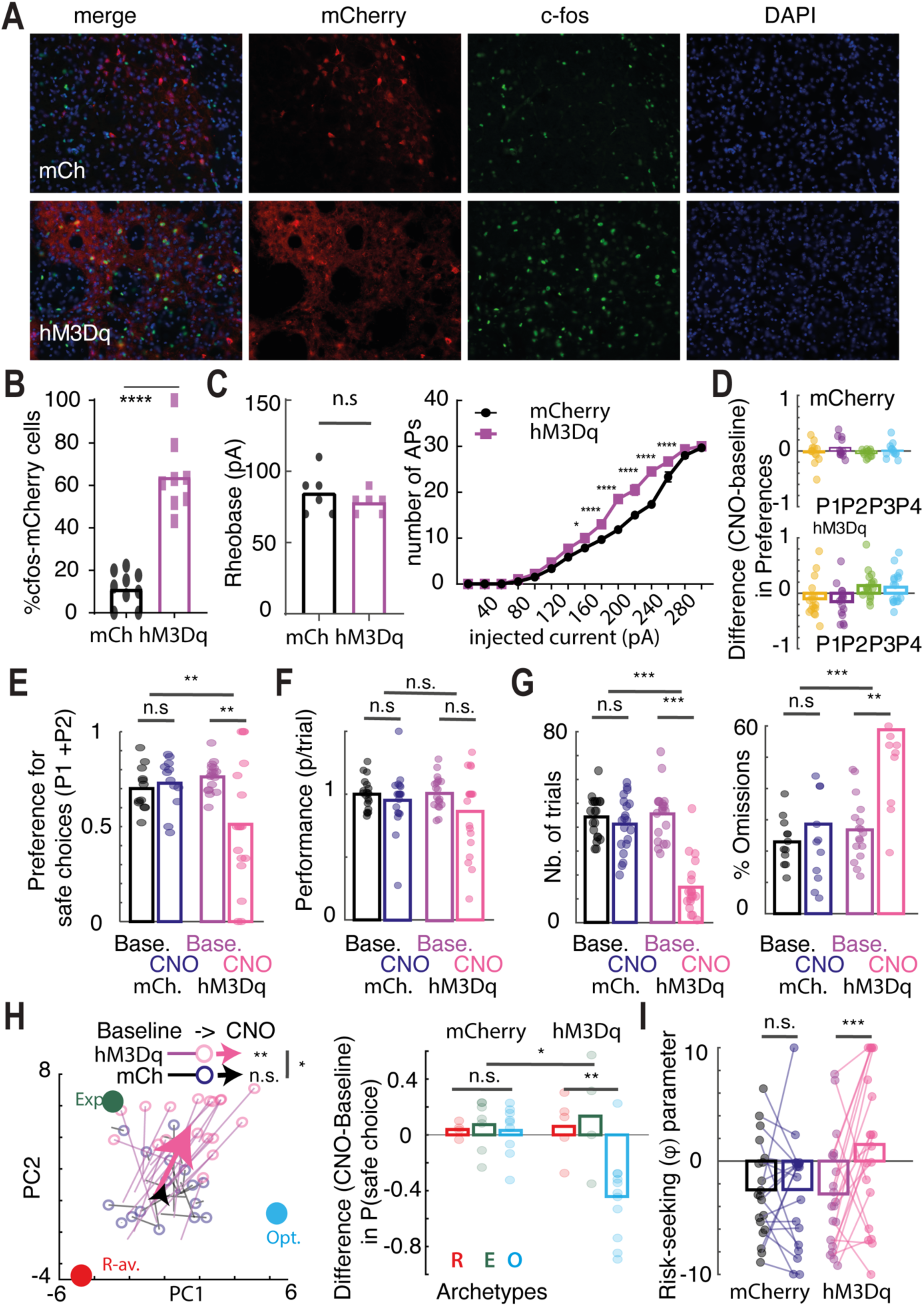
DMS d-SPNs promote risk-seeking. **A.** Microscope fluorescence for control (top) and hM3Dq (bottom) individuals in D1-Cre mice injected in the dorsomedial striatum (DMS): DAPI (blue) indicates the nuclei, mCherry (red) indicates the transfected neurons and Fos (green) indicates the CNO-activated neurons. The arrowheads indicate hM3Dq-expressing neurons. **B**. Fos expression was significantly greater in the hM3Dq group than in the control group. Scale bar= 20 μm. The graph represents the mean±SEM. **C**. Electrophysiological recordings after CNO treatment during the IGT. From left to right: the rheobase did not differ between control and hM3Dq-expressing mice after CNO treatment, but hM3Dq-expressing neurons had increased intrinsic excitability compared with controls. **D.** Differences in all preferences for the four options between CNO and baseline for mCherry controls and hM3Dq mice. **E**. CNO activation of DMS D1-expressing neurons transfected with hM3Dq decreased the preference for the safe options (P1+P2). **F**. CNO treatment did not affect performance (number of pellets divided by the number of trials). **G.** CNO treatment decreased the number of trials and increased the percentage of omissions. **H**. Right: ternary plot showing the global effect of CNO on strategies, with each point (black, mCherry controls; pink, hM3Dq animals) representing an animal, with a connected line showing displacement in the ACP space from baseline to CNO. The larger arrows show the group averages (black, mCherry controls; pink, hM3Dq animals), with hM3Dq animals moving away from the R apex under CNO. Left: data split by archetypes indicate that the decrease in the safe choice was due mainly to an effect on Optimizers. **I**. Data fit by the choice model suggest that the decreased choice of the safe option reflects an increase in risk-seeking upon CNO treatment. n.s. not significant; * p<0.05. ** p<0.01; *** p<0.001; **** p<0.0001.

Having verified that hM3Dq expressed in DMS-dSPNs favored their activation during the mouse gambling test, we next assessed how this increase in neuronal activity affected decision-making. We compared the differences in preferences for the four options (P1-P4) in the mouse gambling task under CNO and at baseline for hM3Dq animals and mCherry controls (Figure 4D). Safe mice (i.e., mice with a preference for the advantageous options P1 and P2) in the hM3Dq + CNO conditions displayed a decrease in the preference for P1 and P2 (Figure 4E, RM-ANOVA, F_(1,38)_=5.83, p = 0.02; CNO effect for the mCherry group: n.s.; for the Gq group: p=0.017). However, this decrease could arise for different reasons: worst decision-making, increased exploration, decreased aversion to punishment, etc. There was no change in overall performance (i.e., average number of pellets per trial; RM-ANOVA, F_(1,38)_=1.48, Figure 4F); hence, we sought a more precise characterization in terms of a modification in decision-making traits. We also observed a decrease in the number of trials and an increase in omissions (Figure 4G; Trials: RM-ANOVA, F_(1,38)_=100, p = 3.10^−12^; CNO effect for the mCherry group: n.s.; for the Gq group: p=2.10^−11^; Omissions: RM-ANOVA, F_(1,38)_=18.08, p = 1.10^−4^; CNO effect for the mCherry group: n.s.; for the Gq group: p=2.10^−6^), i.e., a decrease in the pace of the instrumental action. However, this decrease in decision frequency was not due to motor control, as locomotion was found to increase after DMS-dSPN facilitation (Supp. Figure 5). The combined modification of preferences and instrumental pace suggested a global effect of DMS-dSPNs facilitating archetypal strategies. Indeed, we observed a global shift in the PCA space for hM3Dq animals under CNO, with a displacement away from the risk-averse apex (Figure 4H, RM-ANOVA, F_(1,38)_=5.16, p = 0.03; CNO effect for the mCherry group: n.s.; for the Gq group: p=0.006) that could be observed for all archetypes (Supp. Figure 5). However, the effect on safe choice (P1+P2 choices) was observed only in Optimizer mice (Figure 4H, 2-way ANOVA: F_(1,34)_=5.25, p = 0.01; R+E mice: n.s.; O mice: p=0.005). We thus analyzed the full choice pattern with the computational model. The model-based analysis indicated that the change in the full choice pattern in the hM3Dq-CNO condition was better explained by an increase in the risk sensitivity parameter (RM-ANOVA, F_(1,38)_=4.84, p = 0.03; CNO effect for the mCherry group: n.s.; for the Gq group: p=0.01, Figure 4I, Supp. Figure 5). Overall, the behavioral data and computational analyses suggest that facilitating DMS-dSPN activity decreases the choice of P1 and P2 by increasing proneness to risk.

### Dorsomedial iSPNs favor large gains, exerting strategy-dependent effects

We then compared the above facilitation of DMS-dSPNs with that of DMS-iSPNs (Figure 5A), as these two subpopulations are hypothesized to act in opposition (*20–27*), in synergy (*28*–*33*) or in a complementary manner, supporting a dual selection-suppression function (*34, 56*). In this subpopulation, selective hM3Dq DREADD expression followed by CNO injection during the mouse gambling test effectively increased intrinsic excitability (f-I gain, 2-way ANOVA F_(14,140)_ = 13), driving Fos activation (p<0,0001, U(3)=0, Mann‒Whitney) in targeted neurons (Figure 5B) without any effects on resting membrane potential (t_(8)_=0.39, n.s., rheobase : t_(8)_=0.63, n.s., and AMPA/NMDA ratio: t_(8)_=0.15, n.s., Fig. 5C and Supp. Fig. 6). In contrast to the facilitation of DMS dSPN activity, CNO did not significantly influence preferences in hM3Dq animals compared with mCherry controls when all mice or only safe mice were considered (Figure 5D). However, there was a significant decrease in the performance (pellets/trials) of the animals under DMS-iSPN facilitation (Figure 5E, RM-ANOVA, F_(1,33)_=172.69, p = 1.10^−14^; CNO effect for mCherry group: n.s.; for the Gq group: p=1.10^−7^), suggesting that decision-making had shifted away from reward maximization (due to poorer decision-making or a shift in preferences). Moreover, facilitating the activity of DMS iSPNs with DREADDs decreased the overall number of trials (RM-ANOVA, F_(1,33)_=25.6, p = 8.10^−7^; CNO effect for the mCherry group: n.s.; for the Gq group: p=1.10^−4^) and increased the percentage of omissions (RM-ANOVA, F_(1,33)_=25.08, p = 2.10^−5^; CNO effect for the mCherry group: n.s.; for the Gq group: p=0.001; Figure 5F). We verified in a subset of mice that DMS-iSPNs facilitated decreased locomotion in the operant box (Supp. Figure 6), in contrast with DMS-dSPN manipulation, which was consistent with changes in omissions or in the number of trials independent of locomotor effects. We then looked for an effect on archetypal profiles. Indeed, CNO induced a global shift in the PCA space in hM3Dq animals toward the Explorers (E) apex (Figure 5G, RM-ANOVA, F_(1,33)_=37.69, p = 6.10^−7^; CNO effect for the mCherry group: n.s.; for the Gq group: p=2.10^−4^). The effect of DREADD-mediated facilitation of DMS-iSPN neuronal activity was thus different from that of DMS-dSPNs (i.e., a shift away from the R apex). This led us to examine more closely the effects of CNO on the preferences of hM3Dq animals, depending on their proximity to archetypes rather than on the safe/risky dichotomy. Facilitating DMS-iSPN activity had opposite effects on the preference for safe (P1 and P2) options in Optimizer mice (a decrease in P1+P2 choices) and in the other two archetypes (increased P1+ P2 choices in R+I animals). Hence, the DREADD effect on choices depended on the baseline strategy of the animals. We assessed the state-dependent effects of CNO with the decision model after verifying that the decrease in trial number, observed under CNO (Figure 5F), did not lead to a systematic bias in model parameter estimations, although it introduced more noise in the parameter recovery (Supp. Figure 6). We then characterized which modifications of the decision-making parameters could best account for the differential effect of DMS-iSPN facilitation on choices. We reasoned that a decrease in performance due to poorer decision-making would appear in the model as a decrease in the exploitation (β) parameter, whereas animals could also become less optimal on average because of nonlinear sensitivity to rewards (ρ) or time-outs (T). A modification in the ρ parameter (decrease in the reward saturation with the pellet number) best explained the differential DREADD effect on choices (Figure 5H, Supp. Figure 6, RM-ANOVA, F_(1,33)_=14.23, p = 6.10^−4^; effect of CNO for the mCherry group: n.s.; for the Gq group: p=0.003). Indeed, in reinforcement-learning models such as ours, the alteration of one decision-making parameter can induce opposite behavioral patterns depending on the values of the other parameters (Supp. Figure 6). Here, an increase in reward saturation (decreasing the value of large gains) increased the preference for P1 and P2 when animals were risk averse but decreased the preference for P1 and P2 in risk-neutral mice (Supp. Figure 6). Furthermore, decreases in reward saturation (i.e., preferring 4 pellets over 4 times 1 pellet) deviated the animals from maximizing the average reward, which is consistent with the observed decrease in global performance following DMS-iSPN facilitation. Overall, the computational analyses suggest that facilitating DMS-iSPN activity decreases how the reward saturates with reward size (favoring large gains), exerting a decision profile-dependent effect on choices.

**Figure 5.**
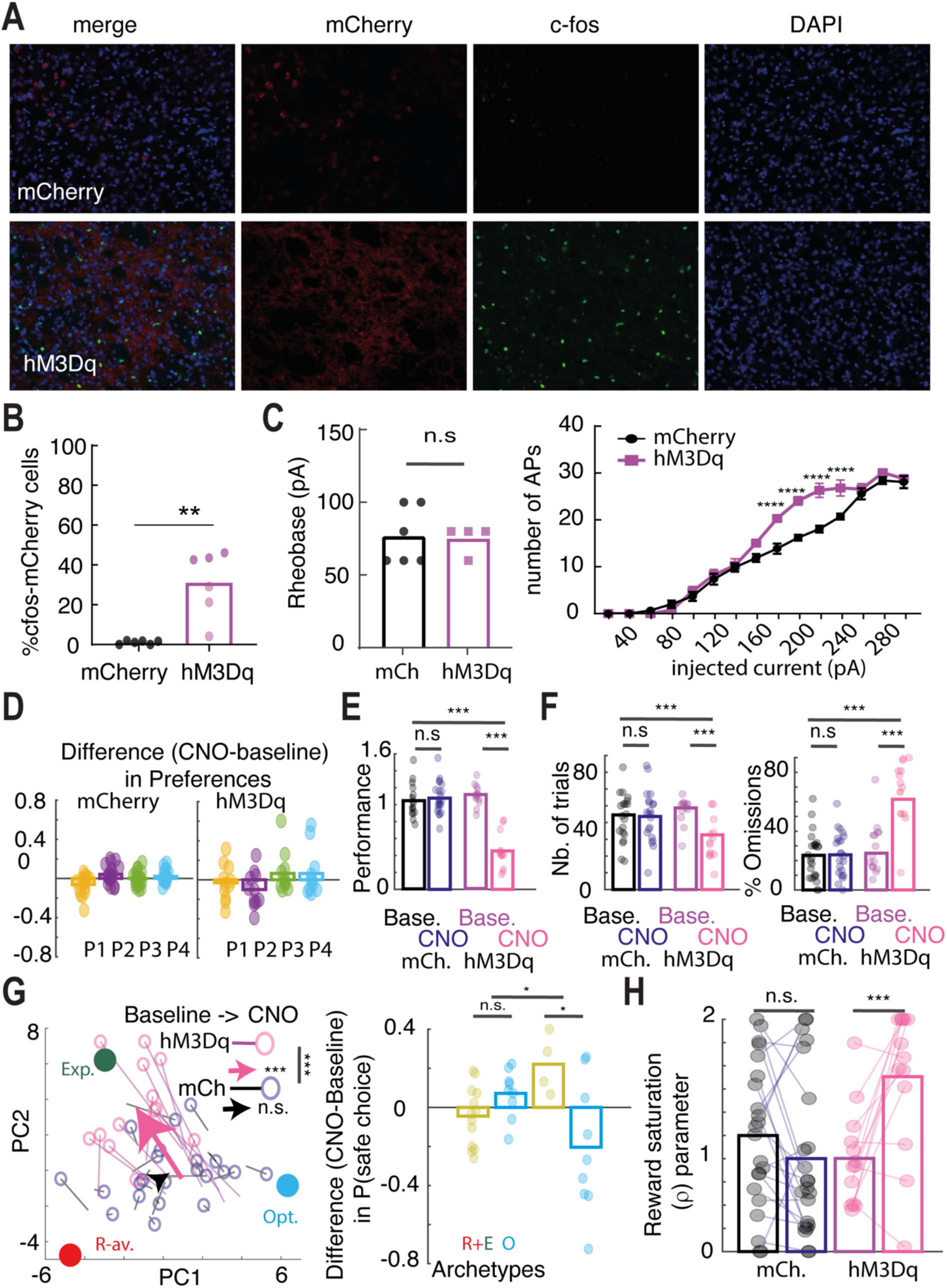
DMS iSPNs favor large gains. **A.** Microscope fluorescence for control (top) and hM3Dq (bottom) individuals in A2A-Cre mice injected in the dorsomedial striatum (DMS): DAPI (blue) indicates the nuclei, mCherry (red) indicates the transfected neurons and Fos (green) indicates the CNO-activated neurons. The white arrowheads indicate hM3Dq-expressing neuronal somas. **B**. Fos expression was significantly greater in the hM3Dq group than in the control group. Scale bar= 20 μm. The graph represents the mean±SEM. **C**. From left to right: the resting membrane potential, rheobase and AMPA/NMDA ratio did not differ between control and hM3Dq-expressing mice after CNO treatment, but hM3Dq-expressing neurons had increased intrinsic excitability compared with controls. **C.** Differences in all preferences for the four options between CNO and baseline for mCherry controls and hM3Dq mice. **E**. CNO treatment decreased performance in hM3Dq animals. **F.** CNO treatment also decreased the percentage of trials and increased the percentage of omissions. **G**. Left: ternary plot showing the global effect of CNO on strategies, with each point (black, mCherry controls; pink, hM3Dq animals) representing an animal receiving CNO treatment, with a connected line showing displacement in the ACP space from baseline to CNO. The larger arrows show the group averages (black, mCherry controls; pink, hM3Dq animals) with hM3Dq moving toward the E apex under CNO. Right: the data split by archetypes indicate the opposite effect of DMS A2A-expressing neurons on the safe choice of Optimizers compared to Risk-averse and Explorers. **H**. Data fit by the choice model suggest that the decreased choice of the safe option reflects a decrease in reward saturation upon CNO treatment. n.s. not significant; * p<0.05; ** p<0.01; *** p<0.001; **** p<0.0001.

### Nucleus accumbens SPNs are less involved in gambling task than DMS

Even if the mouse gambling task is instrumental, the nucleus accumbens (NAc) may also influence choice behavior. Facilitating NAc-dSPN (Figure 6A) activity did not exert effects similar to those of facilitating DMS-dSPN activity. hM3Dq expression coupled with CNO treatment efficiently drove Fos activity and changes in intrinsic excitability in NAc-dSPNs (Figure 6B). In the mouse gambling task, such facilitation of neuronal activity strongly decreased the number of trials (Figure 6C, RM-ANOVA, F_(1,28)_=16.68, p = 3.10^−4^; CNO effect for the mCherry group: n.s.; for the Gq group: p=4.10^−4^), in contrast to DMS-dSPN facilitation, which did not exert any effect. Facilitating NAc-dSPN activity did not alter the proportion of premature or omission trials (Figure 6C), again in contrast with the reduction in these measures following DMS-dSPN manipulation. We did not observe any differences in choice behavior, either following the safe/risky dichotomy or when considering archetypes or overall performance (Supp. Figure 7). However, facilitating NAc-dSPNs increased locomotion (Supp. Figure 7). In the PCA, the overall effect corresponded to a modest (but significant) shift away from the Optimizer (O) apex (Figure 6D, RM-ANOVA, F_(1,28)_=13.53, p = 1.10^−3^; CNO effect for the mCherry group: n.s.; for the Gq group: p=0.03). We did not find any differences in the parameters from the computational model of decision-making when we fitted the choices under the hM3Dq+CNO treatment (Supp. Figure 7). Overall, facilitating NAc-dSPNs had an effect on the decision task, but that was less specific than that of facilitating DMS-dSPNs.

**Figure 6.**
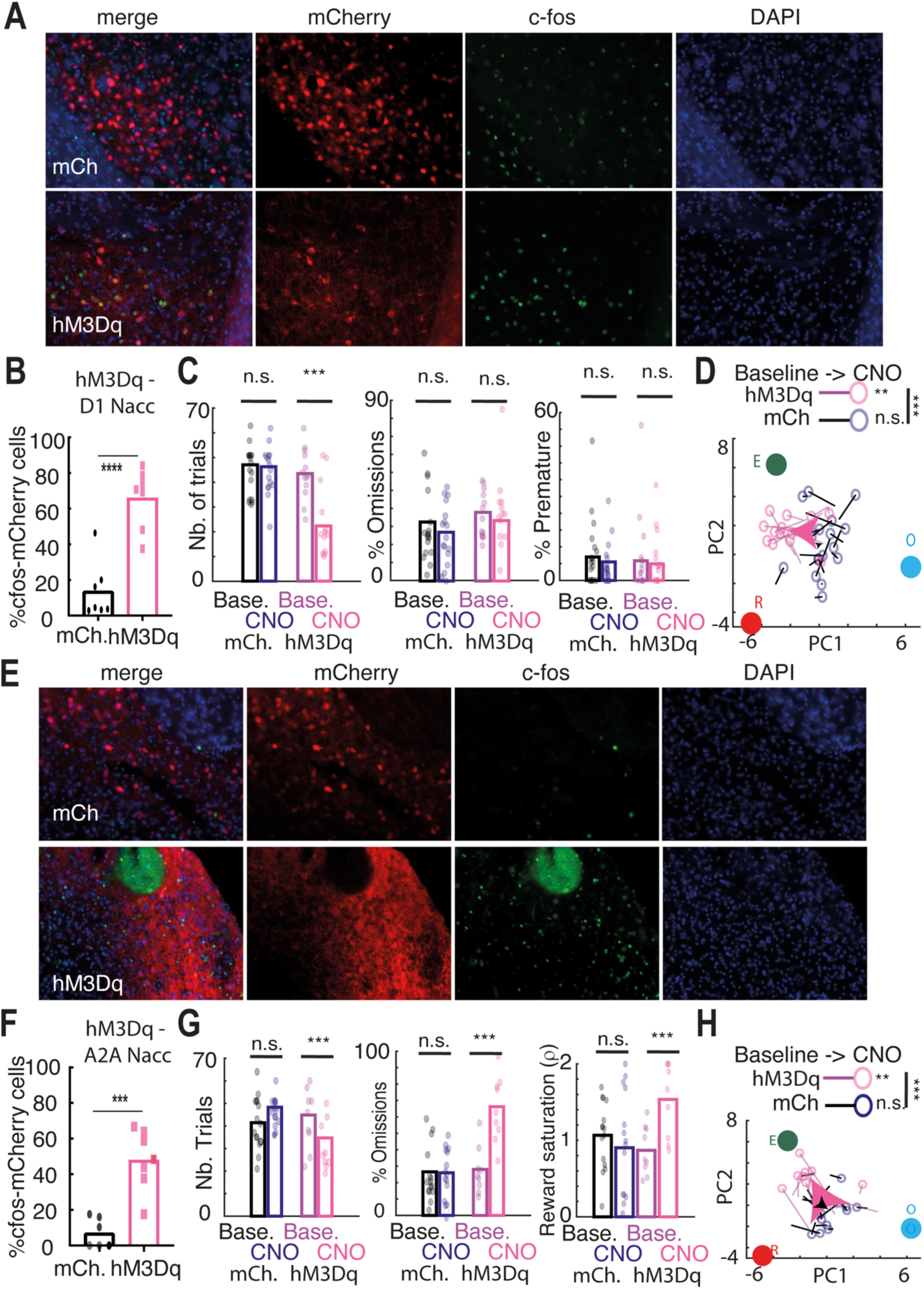
The NAc d- and i-SPNs affect mouse gambling, but less than their DMS counterparts. **A.** Microscope fluorescence for control (top) and hM3Dq (bottom) individuals in D1-Cre mice injected in the nucleus accumbens (NAc): DAPI (blue) indicates the nuclei, mCherry (red) indicates the transfected neurons and Fos (green) indicates the CNO-activated neurons. The white arrowheads indicate hM3Dq-expressing neuronal somas. **B**. Fos expression was significantly greater in the hM3Dq group than in the control group. Scale bar= 20 μm. The graph represents the mean±SEM. **C**. From left to right: CNO treatment decreased the number of trials initiated, increased the percentage of omissions, and decreased the reward saturation parameter from the model fit. **D**. Ternary plot showing the global effect of CNO on strategies, with each point (black, mCherry controls; pink, hM3Dq animals) representing an animal, with a connected line showing displacement in the ACP space from baseline to CNO. The larger arrows show the group averages (black, mCherry controls; pink, hM3Dq animals) with hM3Dq moving toward the E apex under CNO. **E,F.** Same as A but for A2A-Cre mice injected in the NAc. **G**. CNO treatment decreased only the number of trials, but not the percentage of premature responses or the percentage of omissions. **H**. Ternary plot (same as D but for A2A-Cre mice injected in the NAc) with hM3Dq moving away from the O apex under CNO.

By contrast, CNO injection in animals expressing hM3Dq in NAc-iSPNs (Figure 6E) increased intrinsic excitability and drove efficient Fos activation in targeted cells (Figure 6F, p<0,0001, unpaired t test). The behavioral effect of DREADD facilitation on NAc-iSPN activity was relatively similar to what we observed when facilitating DMS-iSPNs. Preferences did not change on average, but the overall number of trials decreased (RM-ANOVA, F_(1,22)_=25.43, p = 5.10^−5^; CNO effect for the mCherry group: n.s.; for the Gq group: p=0.001), whereas the proportion of omissions increased (RM-ANOVA, F_(1,22)_=25.6, p = 8.10^−7^; CNO effect for the mCherry group: n.s.; for the Gq group: p=1.10^−4^, Figure 6G). This pattern was reflected in the PCA space, where CNO induced a shift of hM3Dq animals toward the E apex, albeit smaller than what we observed following DMS-iSPN manipulation (RM-ANOVA, F_(1,22)_=33.22, p = 2.10^−4^; CNO effect for the mCherry group: n.s.; for the Gq group: p=0.005, Figure 6H). Like DMS-iSPNs, facilitation by NAc-iSPNs decreased overall performance and locomotion (Supp. Figure 7). Similarly, the behavioral effects of NAc-iSPN facilitation could be accounted for by an increase in reward saturation in the computational model (RM-ANOVA, F_(1,22)_=26.03, p = 0.03; CNO effect for the mCherry group: n.s.; for the Gq group: p=0.0076, Figure 6G, Supp. Figure 7). Hence, facilitating NAc-iSPNs appeared to exert similar, but smaller in magnitude, effects than facilitating DMS-iSPNs did.

### Both dorsolateral SPN populations have no specific effects on gambling task

Finally, we evaluated the influence of the DLS on choice behavior. CNO treatment drove Fos activity and increased the intrinsic excitability of targeted neurons in hM3Dq animals in the DLS-dSPN (p<0,0001, unpaired t-test, Figures 7A, 7B). We observed small effects on the proportions of omission trials (Figure 7C). However, in the mouse gambling task, there was no change in preferences (in safe mice or when archetypes were considered) under hM3Dq +CNO treatment (Figure 7D). Similarly, CNO treatment drove Fos activity and increased the intrinsic excitability of targeted neurons in hM3Dq animals in the DLS-iSPN (p<0,0001, unpaired t-test, Figure 7E,F). We observed small effects on the proportions of omission trials (Figure 7G), with a decrease in premature trials (Figure 7G) and in locomotion (Supp. Fig 8). As for DLS-dSPN, there was no change in preferences under hM3Dq +CNO treatment (Figure 7H). Consistently, we did not find any significant differences in model parameters when the choices were fitted; under any (DLS-iSPN or DLS-dSPN) conditions (Supp. Figure 8), for both manipulations (DLS-iSPNs and DLS-dSPNs), consistent with the lesser involvement of the DLS in decision-making.

**Figure 7.**
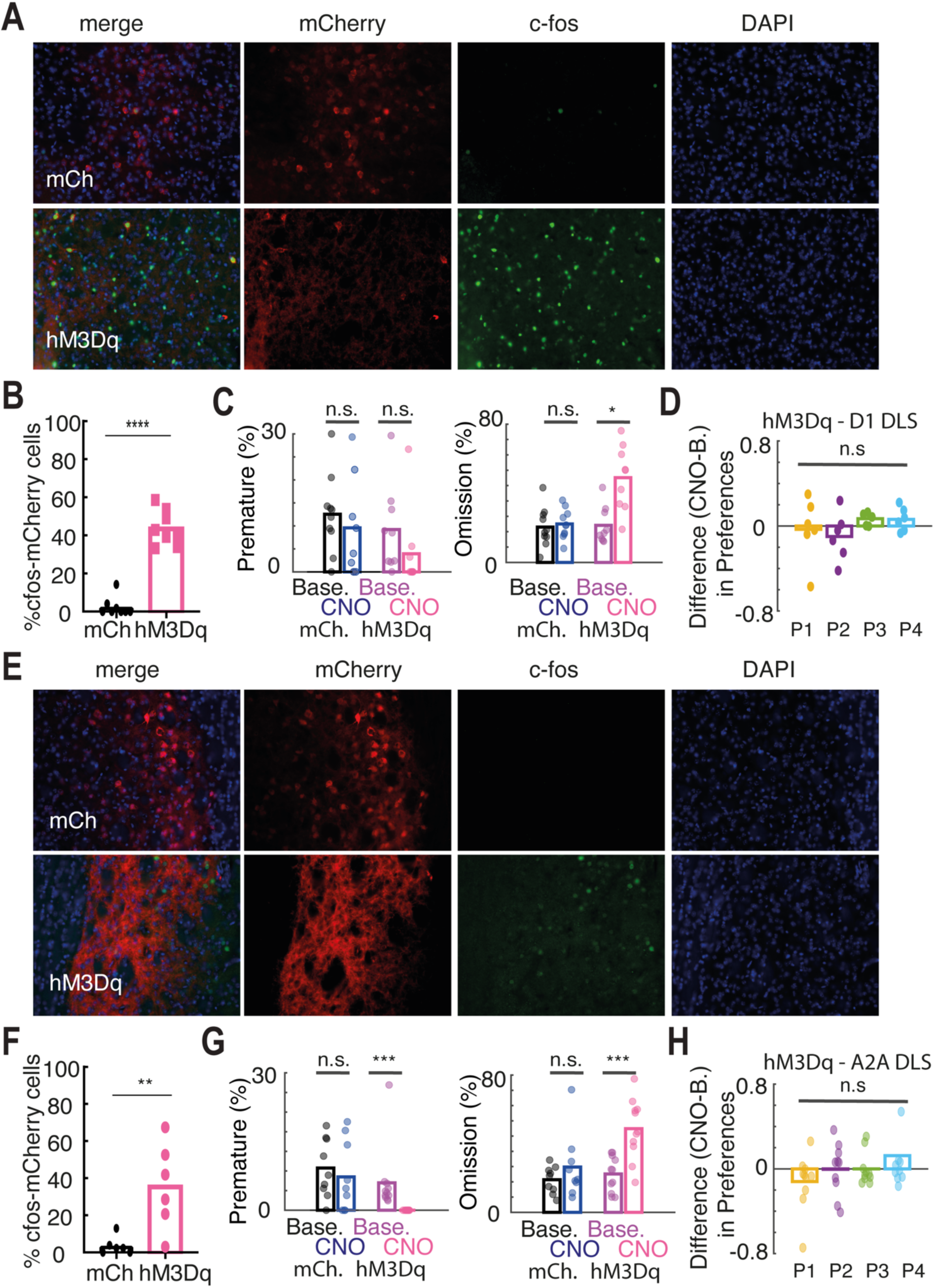
Dorsolateral i-SPN and d-SPN have no specific effects on mouse gambling task. **A.** Microscope fluorescence for control (top) and hM3Dq (bottom) individuals in D1-Cre mice injected in the dorsolateral striatum (DLS): DAPI (blue) indicates the nuclei, mCherry (red) indicates the transfected neurons and Fos (green) indicates the CNO-activated neurons. The white arrowheads indicate hM3Dq-expressing neuronal somas. **B**. Fos expression was significantly greater in the hM3Dq group than in the control group. Scale bar= 20 μm. The graph represents the mean±SEM. **C**. CNO treatment did not affect the proportion of premature response (left), but increased the percentage of omissions (right). **D**. There was no difference in any preference for the four options between the CNO groups and the baseline groups for mCherry controls and hM3Dq DLS-injected D1-Cre mice. **E, F**. Same as A,B for A2A-DLS mice. **G**. CNO treatment decreased the proportion of premature response (left), and increased the percentage of omissions (right). **H**. There was no difference in any preference for the four options between the CNO groups and the baseline groups for mCherry controls and hM3Dq DLS-injected A2A-Cre mice. n.s. not significant; * p<0.05; ** p<0.01; *** p<0.001; **** p<0.0001.

## DISCUSSION

Consistent with other studies in human and rat versions of the IGT, we found marked interindividual variability in preferences. However, we departed from usual classifications in terms of experimenter-defined performance (*37*, *39*). Instead, we chose an unsupervised approach (archetypal analysis) to characterize individuals as intermediate between extrema rather than clearly separated clusters. Whether explicitly or implicitly, assessing interindividual variability on the single scale of “task performance” assumes that (1) animals have evolved to optimize a given quantity, e.g., long-term reward relative to motor costs, and that (2) the experimental setup measures the fitness of the animals on this scale (i.e., the setup is “factor pure”(*57*)). However, organisms have likely evolved to trade priorities across various objectives. In particular, most animals are equipped with a cognitive architecture for deliberative decisions between goals (*58*). Decision-making requires assigning values to options, which is a fundamentally subjective process (*59*, *60*). Computational decision-making models constitute powerful tools designed to infer subjective valuations from the temporal series of choices. However, these models have limitations when applied to experimental data, particularly due to the limited number of choices a rodent can make before reaching satiety, which introduces noise into model parameter estimation (*61, 62*)-especially relevant when interpreting CNO sessions with fewer trials.

Overall, we could describe patterns of interindividual variability in preferences in terms of “cognitive profiles” with animals distributed between extrema (archetypes) rather than strongly defined clusters. Considering several cognitive dimensions rather than a single “adaptive” scale, recasts the “maladaptive” decisions (*50*) concept as “atypical” instead. We further show how cognitive profiles from the archetypal analysis relate to the decision processes in the model (i.e., reward, risk and time-out sensitivities) that generate choices. By moving along these dimensions of decision parameters, animals can exhibit different preference patterns based on their social and physical context, age, and previous experiences (*42*, *63*, *64*). Importantly, as computational models of decision-making are nonlinear, affecting one decision parameter (e.g., reward sensitivity) through chemogenetics may exert effects that depend upon the values of the other parameters. This reinforces the need to infer generative processes from the data rather than focusing purely on overt measures (e.g., performance or a given choice).

Specific manipulations of striatal subpopulations affecting cognitive profiles underscore that decision processes “parameters” are emergent properties from neural interactions. Brain markers correlate with individual variability in choice behavior, such as prefrontal serotonin (*40*) or the balance between striatal and prefrontal excitability (*37*). Nevertheless, much work remains to be done to determine the causality between brain markers and cognitive profiles. Here we detected marked variability in decision-making among a quasiclonal population of inbred mice. Instead of being the origin of interindividual variability in decision-making, brain markers may act as mediators of individual experience and the social context of cognitive strategies (*42*).

To study the causal implications of striatal subpopulations in choice behavior, we preferred DREADD over optogenetics because a temporal window for manipulation was not needed. Additionally, decision-making likely occurs during the whole task. Furthermore, we wanted to avoid the risk of nonphysiological synchronization of striatal neurons under specific optogenetic stimulation protocols. Indeed, striatal neurons are involved in action selection (*29*, *65–68*), and neurons promoting distinct goals (or weighting different decision parameters) are not activated synchronously (*30*, *33*, *34*). DREADDS partially circumvents this issue as a gain-of-function approach in this study. However, one limitation of our DREADD approach is that the CNO ligand may be converted to clozapine (*69*), an antipsychotic with sedative effects potentially decreasing behavioral impulsivity (*70–72*), but our mCherry controls excluded DREADD-independent effects.

Our results precise and extend the literature on the differential involvement of subparts of the striatum (*73*). The role of the DMS in sustaining instrumental associations (action-reward) has been widely proven before (*6*, *7*, *13*, *16*, *74*). Here, we show that the DMS is critical not only for maintaining the instrumental response of mice but also for determining their preferences under risk. We further provide a computational rationale (and iSPN versus dSPN distinction) for the effect of the DMS on risky choices (*75*) in terms of sensitivity to reward variance and reward saturation. While the NAc has also been implicated in risky decision-making, with specific involvement of iSPNs, as in our study(*15*), we found a lower NAc effect than the that of the DMS. This may be due predominantly to the instrumental nature of the task, as the NAc supports the acquisition of stimulus‒outcome associations (*76–79*). There was no effect of DLS manipulations in the IGT, which is consistent with the known dissociation of the DMS and DLS (*12*, *13*, *17*) in controlling goal-directed behavior and habits, respectively. Despite extensive training, the mice did not develop rigid preferences (i.e., most mice had balanced choice behavior; Figure 1), suggesting that they did not develop habits (although we did not test choices in extinction).

We also provide novel data on the major anatomical and functional distinctions in the basal ganglia between the direct (dSPNs) and indirect (iSPNs) pathways. Concurrent views have proposed that dSPNs and iSPNs may work either in opposite or complementary ways to promote and oppose (or refine) action selection (*33*, *64*, *80–82*). However, in the DMS, we found that DREADD-mediated activation of both iSPNs and dSPNs enhanced risky choice through distinct computational mechanisms (i.e., risk sensitivity and reward saturation, respectively). This points to more complex interactions between pathways. The activity of iSPNs has been related to the encoding of nonrewarding events or changes in reward value (prediction errors), leading to the updating of action value (*36*) and henceforth task switching (*82*, *83*). Our computational account is consistent with this view, as reward saturation (i.e., how the subjective value saturates with reward magnitude) may arise from learning effects. However, we could not test this hypothesis directly, as including a learning rate induced correlations with explore/exploit and risk-seeking parameters and led to poor parameter recovery, probably due to the relatively small number of trials in the mice. As we could disentangle reward saturation from sensitivity to uncertainty in our multiple-choice setup, our results extend and explain those obtained in tasks involving binary choices. For example, activating iSPNs may increase seemingly stochastic choices in serial choice tasks in which the exploratory choice may also constitute a risky (high variance) choice (*84*) because reward saturation affects the perception of risk (Supp. Figure 5).

Our findings reframe previous results on dopamine manipulation. DAT-knockdown mice make riskier choices in the IGT(*49*), and “safe” decision-makers show lower dopamine levels in the dorsal striatum. This suggests dopamine’s effects on risk-taking arise from distinct D1R- and D2R-mediated mechanisms. The next step will be to use correlative approaches such as in vivo calcium imaging to characterize the physiological responses of DMS-dSPNs during choice preference. Indeed, concerted and cooperative activity between both striatal pathways is needed for proper action initiation and execution (*28–34*, 56, *66*). It will then be possible to compare how decisions are encoded in the different types of SPNs and compare such encoding between the Risk-averse, Explorers, and Optimizers mice. In addition, a correlation between the variables of the computational model and the activity patterns of certain dSPNs or iSPNs can also be found (*85*). Moreover, we could also measure whether, as our results suggest, the activity of dSPNs when mice are making a choice differs among the DMS, DLS and NAc. Using in vivo techniques, our results suggest that increasing neuronal excitability rather than directly activating targeted cells is beneficial. These questions should be carefully investigated in future work.

Understanding and aiding individuals in the context of decision-related disorders, such as pathological gambling (*86*) and drug addiction (*87*), necessitates a shift in perspective. Impairments in decision-making should be seen as discrepancies between anticipated outcomes and actual choices. We propose a framework where decision-making “profiles” arise from evolutionary trade-offs, shaped by parameters from the deliberative machinery leading to goal-directed choice. Biological and social (*42*) factors can shift the spectrum of potential strategies, likely governed by “meta-learning” rules (*88*), offering promising avenues for translational research.

## METHODS

### Animal care and use

All procedures were performed according to the Institutional Animal Care Committee guidelines and were approved by the Local Ethical Committee (Comité d’Ethique et de Bien-Être Animal du pôle santé de l’Université Libre de Bruxelles (ULB), Ref. No. 646 N). The mice were maintained on a 12-hour dark/light cycle (lights on at 8 pm) with *ad libitum* access to food and water. The room temperature was set to 22 ± 2 °C with constant humidity (40–60%). The behavioral tests were performed during the dark photoperiod. Both male and female transgenic mice (≥ 8 weeks) were used in all the behavioral experiments.

### Generation of transgenic mice

The genetic background of all the transgenic mice used in this study was C57BL/6J. The mice were heterozygous and maintained by breeding with C57BL/6 mice. Two transgenic mouse lines were used: A_2A_-Cre (*22*) and D_1_-Cre (EY262; GENSAT)(*54*). Simple transgenic A_2A_-Cre or D_1_-Cre mice were used for the virus-mediated targeting of iSPNs or dSPNs, respectively.

### Viral injections

Under Avertin anesthesia (2,2,2-tribromoethaol 1.25%, 2-methyl-2-butanol 0.78%; 20 µL/g, i.p.; Sigma Aldrich), male A_2A_-Cre and D_1_-Cre mice (≥ 8 weeks old), which allowed us to target A2A- and D1-expressing neurons, respectively, received two injections (at 100 nL/min) under stereotaxic control in the DLS (AP +0,8 L ±2,42 DV −3,2), DMS (AP +1,2 L ±1,33 DV −3,2) or NAc (AP +1,95 L ±1,2 DV −4,85) of a Cre-dependent virus encoding hM3Dq (AAV5-hSyn1-DIO-hM3Dq-mCherry, Addgene) or mCherry alone as a control (AAV5-hSyn1-DIO-mCherry, Addgene). The coordinates were relative to Bregma according to the Franklin and Paxinos atlas third edition. The injection volumes were as follows: DLS 0.4 μl, DMS 0.35 μl, and NAc 0.2 μl. The injections were delivered through a cannula connected to a Hamilton syringe (10 µL) placed in a syringe pump (KDS-310-PLUS, KDScientific). The cannulas were lowered into the brain and left in place for 10 min after infusion. A minimum of 3 weeks elapsed between the stereotaxic injections to ensure optimal protein expression levels. For all the mice, the accuracy of the injections was checked under a microscope after behavioral testing. The transfected area could be identified via mCherry staining (see Supp. Figure 4). Behavioral data from animals whose targeted area was not accurate or unilaterally injected were excluded from the analyses (see table below).

| Mouse strain | Striatal area | Injected virus | Total n° of mice injected | **Excluded**(bad stereotaxic targeting) |
| --- | --- | --- | --- | --- |
| D<sub>1</sub>-Cre | DMS | mCherry | 20 | 0 |
|  |  | hM3Dq | 21 | 1 |
|  | DLS | mCherry | 10 | 0 |
|  |  | hM3Dq | 10 | 1 |
|  | NAc | mCherry | 23 | 6 |
|  |  | hM3Dq | 20 | 7 |
| A<sub>2A</sub>-Cre | DMS | mCherry | 22 | 0 |
|  |  | hM3Dq | 15 | 2 |
|  | DLS | mCherry | 9 | 0 |
|  |  | hM3Dq | 11 | 0 |
|  | NAc | mCherry | 15 | 0 |
|  |  | hM3Dq | 12 | 3 |

### Mouse gambling task

Behavioral testing was performed in 6 identical operant chambers (IMETRONIC, Pessac, France). Four circular holes were available in a curved wall (the central hole was obstructed, and only 4 holes were actually available), with LED lights at the rear of the chamber and a food dispenser of 20 mg chocolate pellets (purified dustless precision pellets for rodents, Bioserv, New Jersey, USA) on the opposite wall. The infrared beams in the nose-poke holes and the magazine allowed the detection of entries to the nose-poke holes. Additionally, a house light located in the ceiling of the chamber was used under specific conditions. These chambers were in individual soundproof closets to avoid any external disturbances during training. Video cameras placed on the top of the chambers allowed tracking of every session. The chambers were connected to a computer and controlled by POLY software (IMETRONIC).

We used a rodent version of the Iowa Gambling Task (IGT) adapted from Young and colleagues (2011)(*49*). Mice have to choose among 4 different options associated with a certain gain and a certain loss (Figure 1). To achieve that goal, mice must undergo several phases of training. The mice were placed on a reduced diet two days before training, aiming to reach 85–90% of their original weight to enhance performance, given that food motivation drives the IGT. Initially, they received 2.5 grams of food per mouse, which was adjusted the next day on the basis of individual weight loss. During IGT training, food amounts remained stable but could be adjusted if the mice fell out of the 85–90% weight range. Once the final training phase commenced, the food quantities were fixed to avoid performance interference. The whole protocol lasted between 1 month and a half and 3 months, depending on the animal. The mice were trained daily at 9 am in a reverse light/dark cycle.

#### Magazine habituation

The mice were trained for 10 minutes per day for 3 consecutive days. During these sessions, every light was turned off. Fifteen seconds after the start of the session, the pellet dispenser was activated to distribute a single pellet. The LED lights of the magazine were then lit. The lights were turned off when the mice obtained the pellet, and the 15 second cycle started again. The magazine would stay lit until retrieval of the pellet to allow an association between the lit magazine and food to be formed.

#### Nose‒poke habituation

Four nose-pokes holes were available, and a lit nose-poke meant that it was an active nose-poke. This training was divided into two phases. In the initial stage, which lasted 4 days, the mice responded to one active nose-poke out of 4, which was chosen randomly in two daily sessions. If successful, 3 pellets were dispensed; otherwise, the sessions could last as long as 10 minutes. For the first 2 days, the pellets were placed in active nose-pokes holes to encourage exploration. In the second phase, all 4 nose-pokes holes remained active throughout the 30-minute sessions, delivering a single pellet upon visitation. Training was continued until the mice achieved 40 responses in two consecutive days, with individual progress varying. Upon reaching this criterion, the mice progressed to the next training phase.

#### Forced-choice IGT

Different nose-pokes holes had specific rewards and punishments. The mice underwent 3 days of forced 30-minute sessions, where a random nose-poke hole was activated, leading to either a reward or punishment afterward.

#### Final IGT

The mice underwent 30-minute sessions with up to 100 trials. Sessions began with a lit magazine that the mice had to approach. A 5-second interval preceded the illumination of 4 nose-poke hole lights; premature responses triggered a time-out. After the interval, mice had 10 seconds to complete a nose poke, and omissions led to new trials. Correct responses extinguished lights and rewarded mice on the basis of specific nose-poke hole probabilities (P1, P2, P3, and P4). Rewards involved pellet delivery, followed by an 8-second period before the next trial began. Punishments initiated a time-out with flashing lights, after which the mice could start a new trial. Perseverative responses were noted but not punished. The mice were trained until their preference stabilized over four consecutive days, establishing a baseline.

### CNO treatment

When the mice reached a stable pattern of preference over 4 days of IGT (see below), we started treatment with 0.9% NaCl for 2 days. All injections were applied 30 minutes before the start of the session. After receiving the CNO injection (1 mg/kg), the mice underwent their last session of IGT. Immediately after the end of this session, the mice were euthanized. A part of the group was used to perform the electrophysiological recordings. In this group, there was a balanced representation of hM3Dq-expressing mice and control mice.

### Immunohistochemistry

After the behavioral experiments were completed, the mice were deeply anesthetized with avertin (2,2,2-tribromoethaol 1.25%, 2-methyl-2-butanol 0.78%; 20 µL/g, i.p.; Sigma Aldrich) and transcardially perfused with PBS followed by 4% paraformaldehyde in PBS. The brains were removed and postfixed overnight at 4 °C. Then, 30-µm coronal slices containing the striatum were cut with a vibratome (VT1000 S; Leica) and stored in PBS. The sections were incubated overnight with a dilution (1/2000 in 1% NHS 0.1% PBST) of a rabbit Fos primary antibody (Santa Cruz). Next, the slices were washed twice for 5 min with PBST and incubated for 1.5 h with a dilution (1/200 in 1% NHS 0.1% PBST) of donkey anti-rabbit secondary antibody Alexa 647 (far red) (Jackson ImmunoResearch). The slices were then washed twice for 5 minutes with PBST. Finally, DAPI nuclear staining was performed (10 minutes in a 1/5000 DAPI solution in 0.01 M PBS), and the samples were washed for 5 min at 0.01 M PBS, mounted on glass slides and cover slipped with Fluoromount.

For the GFP immunostaining, the protocol was the same as that for the chicken anti-GFP primary antibody (1/2000, Santa Cruz) and the goat anti-chicken secondary antibody Alexa 647 (1/400, Jackson ImmunoResearch).

Image acquisition was performed with a Zeiss Axioimager Z1 at 20X magnification, and the images were processed with AxioVision (Zeiss) software. Cell counting was performed with the open-source software FIJI. Additionally, we used Axio Zoom.V16 (Zeiss) and its tiling tool to obtain whole-slice surface images.

#### Acute brain slice preparation

Recordings were made ex vivo on brain slices that were kept alive. The animals were anesthetized with halothane and then euthanized via decapitation. Their brains were quickly collected, adhered to a vibratome plate and submerged in a vibratome container filled with cutting solution at 4 °C. Brain slices were cut (220 μm thick) and then transferred to an incubator chamber filled with artificial cerebrospinal fluid (aCSF) (Table MM-1) at 34 °C for at least 60 minutes before the recordings started. This resting period is required for the recovery of proper neuronal metabolic activity. Both the cutting solution and aCSF were constantly oxygenated, as they were supplied with a 95% O_2_ and 5% CO_2_ mixture to avoid hypoxia. After recovery, the brain slices were transferred one by one to the recording chamber, which was constantly perfused with oxygenized aCSF at room temperature. The setup was equipped with a camera and a Zeiss upright microscope (Axioskop 2FS Plus, Zeiss), which first allowed for the identification of different areas in the striatum in the slices (5x/0,15 EC PlanNEOFLUAR objective lens, Zeiss) and secondly, for the identification of different cell populations (63x water-immersion objective, Zeiss). Depending on the group of mice tested, we recorded from either the DLS, DMS or NAc (according to the injection site of the group).

#### Recordings

The patch pipette was obtained with a vertical two-stage puller (PIP 5, HEKA), and during the recordings, it was filled with intracellular solution (Table MM-1). The Ag/AgCl electrode in the pipette was connected to an EPC-10 (Heka) amplifier, whose signal was recorded with Patchmaster software (Heka). Intrinsic excitability was studied via a current clamp. First, the resting membrane potential was measured (without applying any current). Afterward, a negative amount of current was injected into the cell to maintain its membrane potential at a value of −80 mV. Increasing intensity currents (increase of 10 pA) were then injected to depolarize the cell and trigger action potentials. Rheobase is defined as the minimum amount of injected current that allows the induction of an action potential. Evoked excitatory postsynaptic currents (EPSCs) were recorded with the internal solution supplemented with spermine (0.1 mM). Spermine blocks calcium-permeable AMPA-Rs at positive potentials and thus allows us to distinguish calcium-permeable from noncalcium-permeable AMPA-R currents(*89*). Evoked EPSCs were isolated from GABAergic currents via the application of gabazine (or SR-95531, 10 μm). We do not make any reference to kainate-Rs because of the difficulties in distinguishing them from AMPA-Rs, as they cannot be easily separated pharmacologically(*90*). Thus, we will use shortcut AMPA-R currents to refer to EPSCs resulting from both AMPA-Rs and kainate-Rs. Evoked EPSCs were recorded in the striatum (DLS, DMS or NAc) by placing a bipolar stimulating electrode (SNEX-200, Science Products GmbH) in the corpus callosum (white matter between the cortex and the striatum) to ensure specific stimulation of the corticostriatal pathway. The duration of the stimulation current pulses was constant and set at 0.2 ms. The intensity of the stimulation was adjusted for each cell and set at the minimal value needed to evoke the largest eEPSC (from 600 to 1000 μA). The slices were stimulated every 10 seconds. Evoked EPSCs were recorded successively at −70 mV, 0 mV and +40 mV to allow computation of the ratio between AMPA-R-mediated currents and NMDA-R-mediated currents. The AMPA-R/NMDA-R ratio provides a sensitive measure to detect differences in glutamatergic synaptic strength between two experimental groups(*91*). AMPA-R-mediated EPSCs were obtained at a holding potential of −70 mV, where no NMDA-R conductance was observed, due to the magnesium block. The sum of AMPA-R- and NMDA-R-mediated EPSCs was evoked at a holding potential of +40 mV. In MSNs, AMPA-R-mediated currents were pharmacologically isolated via the application of L689,560 (10 μM), a noncompetitive NMDA-R antagonist, to the solution. The NMDA-R current was obtained via off-line subtraction of the two traces. eEPSCs were digitized at a frequency of 10 kHz and filtered online with a low-pass Bessel filter, using a cutoff frequency set at 3 kHz. Ten eEPSC traces were averaged for each condition.

### Statistical analyses

#### General statistical analyses

The results are plotted as the means ± sems. The total number (n) of observations in each group and the statistics used are indicated in the figure legends. Classical comparisons between means were performed via parametric tests (Student’s t test or ANOVA for comparing more than two groups when the parameters followed a normal distribution (Shapiro test, P > 0.05) and nonparametric tests (here, Wilcoxon or Mann‒Whitney tests) when the distribution was skewed. Multiple comparisons were corrected via a sequentially rejective multiple test procedure (Holm). Probability distributions were compared via the Kolmogorov– Smirnov (KS) test, and proportions were evaluated via the chi-square test (χ2). All the statistical tests were two-sided. P > 0.05 was considered not to indicate statistical significance.

#### Archetypal analysis

Computations were performed via the ParTI routine in MATLAB (*53*). Briefly, given an n × m matrix representing a multivariate dataset with n observations (n = number of animals) and m attributes (here, m = 7, the 4 preferences, the number of trials, the percentage of premature responses and the percentage of omissions), the archetypal analysis finds the matrix Z of k multidimensional archetypes (k is the number of archetypes). k was forced to be lower than m so that there cannot be more groups than dimensions. Z is obtained by minimizing || X-α ZT | | 2, with α representing the coefficients of the archetypes (αi,1…k ≥ 0 and ∑ αi,1…k = 1), and | |.||2 representing a matrix norm. The archetype is also a convex combination of the data points Z = XTδ, with δ ≥ 0, and their sum must be 1. The α coefficient depicts the relative archetypal composition of a given observation. For k = 3 archetypes and an observation i, αi,1; αi,2; αi,3 ≥ 0 and αi,1 + αi,2 + αi,3 = 1. A ternary plot can then be used to visualize the data (αi,1; αi,2; αi,2) used to assign an individual behavior to its nearest archetype (i.e., k max(αi,1; αi,2; αi,3)). αi,j are also used as variables to estimate population archetypal composition. The pure archetype corresponds to 1, the archetypal composition decreases linearly with increasing distance from the archetype, and 0 corresponds to points on the opposite side.

### Reinforcement-learning model

#### Decision model

Decision-making models describe the probability Pi of choosing the next state i as a function (the “choice rule”) of a decision variable. We modeled decisions between the four options with a “softmax” decision rule, defined by P_i_ = e−ß(sum(*V*_a_)/(1 + exp(−ß(sum(*V*_i_))), where β is an inverse temperature parameter reflecting the sensitivity of choice to the difference of decision variables (values) *V*_i_. A large β corresponds to exploitation, i.e., choosing the option that seems the best thus far, whereas a small β corresponds to exploration (at β=0, all options are chosen equally).

#### Decision variable

The decision variable or value *V* of an option is modeled as *V* = *R* + *P* + *U*, i.e., the sum of the expected (average) reward, the expected punishment, and the expected uncertainty (risk). The expected reward was given by *R* = *p*\*(Number of Pellets)^(ρ), where *p* is the probability that the choice is rewarded, and ρ is a model parameter of how the subjective value of the option depends on the reward magnitude. A ρ value close to 1 corresponds to a linear subjective value (the subjective value equals the reward size). A value of ρ smaller than 1 corresponds to reward saturation, e.g., the subjective value of 4 pellets is less than 4 times the value of one pellet. The expected punishment depends on the opportunity cost of time *Tc*, i.e., the average reward to be lost if there is a time out: *P* = (1-*p)*\*(TimeOut duration)^(*T*)*\*Tc;* where *T* scales the perception of the time-out duration. A *T* value close to 1 corresponds to a linear perception of time, and a γ smaller (larger) than 1 corresponds to underweighting (respectively, overweighting) long durations. Risk corresponded to variance in the outcome, *V*(*X*) = *E*(*X*^2^) − *E*(*X*)^2^= *pR*^2^ − (1 − *p*)*P*^2^ − (*pR* − (1 − *p*)*P*)^2^ ; scaled by the free parameter φ (a negative φ corresponds to risk aversion, and a positive φ corresponds to risk seeking).

#### Fitting the model

The free parameters of the model (β, ρ, *T*, φ) were fitted by maximizing the data likelihood. Given a sequence of choices c = c1…T, the data likelihood is the product of their probability given by the softmax decision rule. We used the fmincon function in MATLAB to perform the model fitting, with the constraints that β ∈]0,10], φ ∈ [−10,10] and *Rs* ∈]0,2] and *Rp* ∈]0,2].

#### Generative properties of the model

To simulate the choice patterns for the different archetypes, we simply displayed the choice probabilities (toward which a simulation would converge at a large n) obtained for the average set of parameters fitted on the average of the individuals assigned to an archetype.

#### Recovery analysis

N_animals_ series of n_trials_ choices were generated for a set of parameters (β, ρ, *T*, φ) and then fitted with the above procedure to assess the quality of parameter recovery. In Supp. Figure 2, we used systematic variations of one parameter with the others held constant close to average fit from the data (β_0_ = 2, ρ_0_=0.8, *T*_0_=1, φ_0_=-2), and N_animals_ =20; n_trials_=50. For Supp. Figure 5, we performed N_animals_ =200 pairs of simulations, one under a control n_trials_ =50 and one under CNO n_trials_ =20, each pair with the same set of parameters drawn from a normal distribution around the average parameters fitted from the data (β_0_ = 2, ρ_0_=0.8, *T*_0_=1, φ_0_=-2). We then fitted the model parameters for each condition separately.

#### Model comparison

We used the Bayesian information criterion (twice the negative log likelihood plus the number of parameters times the log of the number of trials) to test whether simpler versions of the model could provide more parsimonious fits. We also displayed some of the model parameter fits in simpler models with a divergence toward limit values to show poorer fits.

## Acknowledgments

We thank Christophe Varin for their critical reading of the manuscript and Philippe Faure for the discussions on the archetype analysis. We thank Delphine Houtteman, Souad Laghmari, Perrine Hagué and Laetitia Cuvelier for their technical assistance and the mouse colonies in Brussels, and LIMIF for their assistance with the imaging techniques. ECR was a research fellow of FRS-FNRS-FRIA (Belgium). JN is a researcher of the CNRS, AKE is a Research Director of the FRS-FNRS and a Welbio investigator

## Author contributions

Conceptualization: ECR and AKE; methodology and investigation for the in vivo and ex vivo experiments: ECR and DR; methodology and investigation for the archetypal analysis; and reinforcement-learning model: JN; formal analysis: ECR, JN, DR; validation and supervision: AKE; writing: ECR, JN and AKE.

## Data and materials availability

All data are available in the manuscript or the supplementary materials.

## List of supplementary materials

**Supplementary Figure 1.**
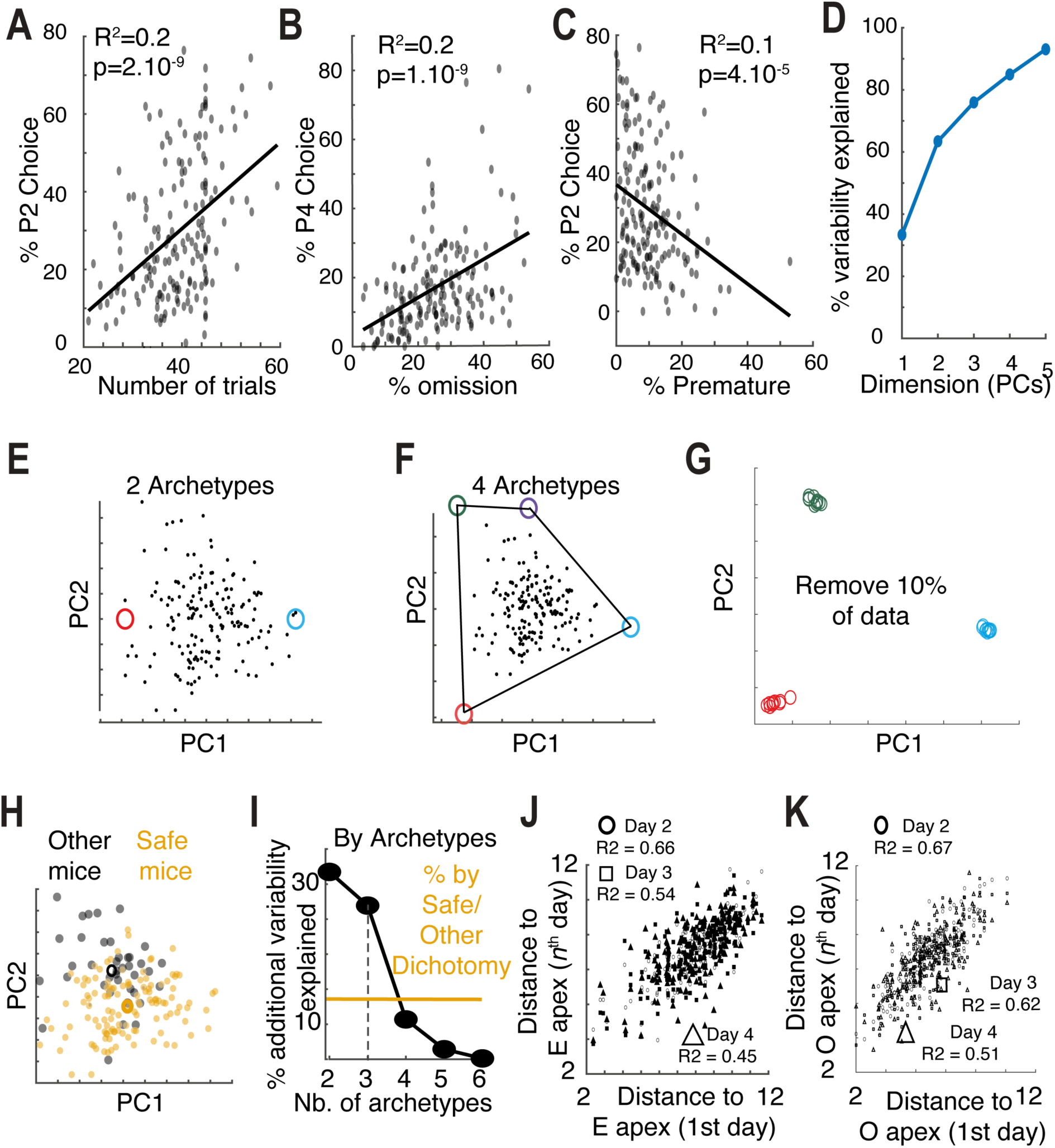
Other examples of correlations between behavioral measures. **A**. Preference for the P2 choice was positively correlated with the number of trials (R^2^ = 0.2, p=2.10^−9^). **B**. Preference for the P4 choice was positively correlated with % of omissions (R^2^ = 0.2, p=1.10^−9^). **C**. Preference for the P2 choice was negatively correlated with % of premature responses (R^2^ = 0.2, p=4.10^−5^). (**D**) % of variance explained by the principal components. (**E,F**) Apices corresponding to 2 archetypes (**E**) and 4 archetypes (**F**). **G**. Apices found when 10% of the data was randomly removed. (**H**) Projection in the PC (1^st^ and 2^nd^) space of mice according to the safe/not safe classical dichotomy. (**I**) Comparison of the % of explained variance by archetypes (black, same as Fig. 2C) and by safe/not safe dichotomy (yellow). (**J,K**) Correlation over days between distances (same as Fig. 2D) to the E (**J**) and O (**K**) apices.

**Supplementary Figure 2.**
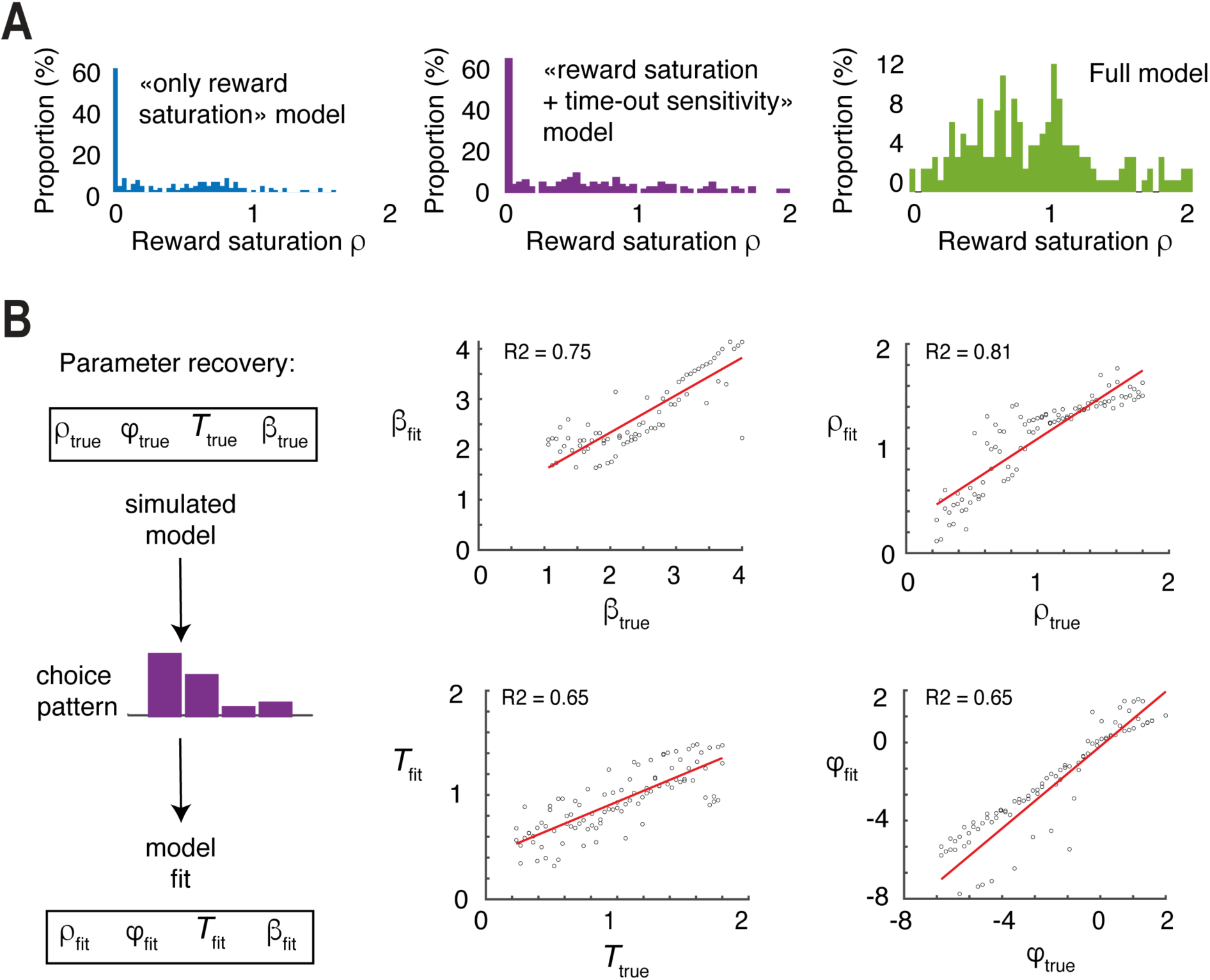
Model comparisons. **A**. Distribution of fitted values for reward saturation parameter in simple model versus the full model; showing that simpler versions of the model fit the data with extreme, implausible values for the parameters (i.e. reward saturation is expected to lie to close to 1 or slightly below 1). **B**. Parameter recovery: choices were generated with multiple realization of a model with controlled parameter (“true” parameters) and these choice patterns were fitted with the model (“fit” parameters).

**Supplementary Figure 3.**
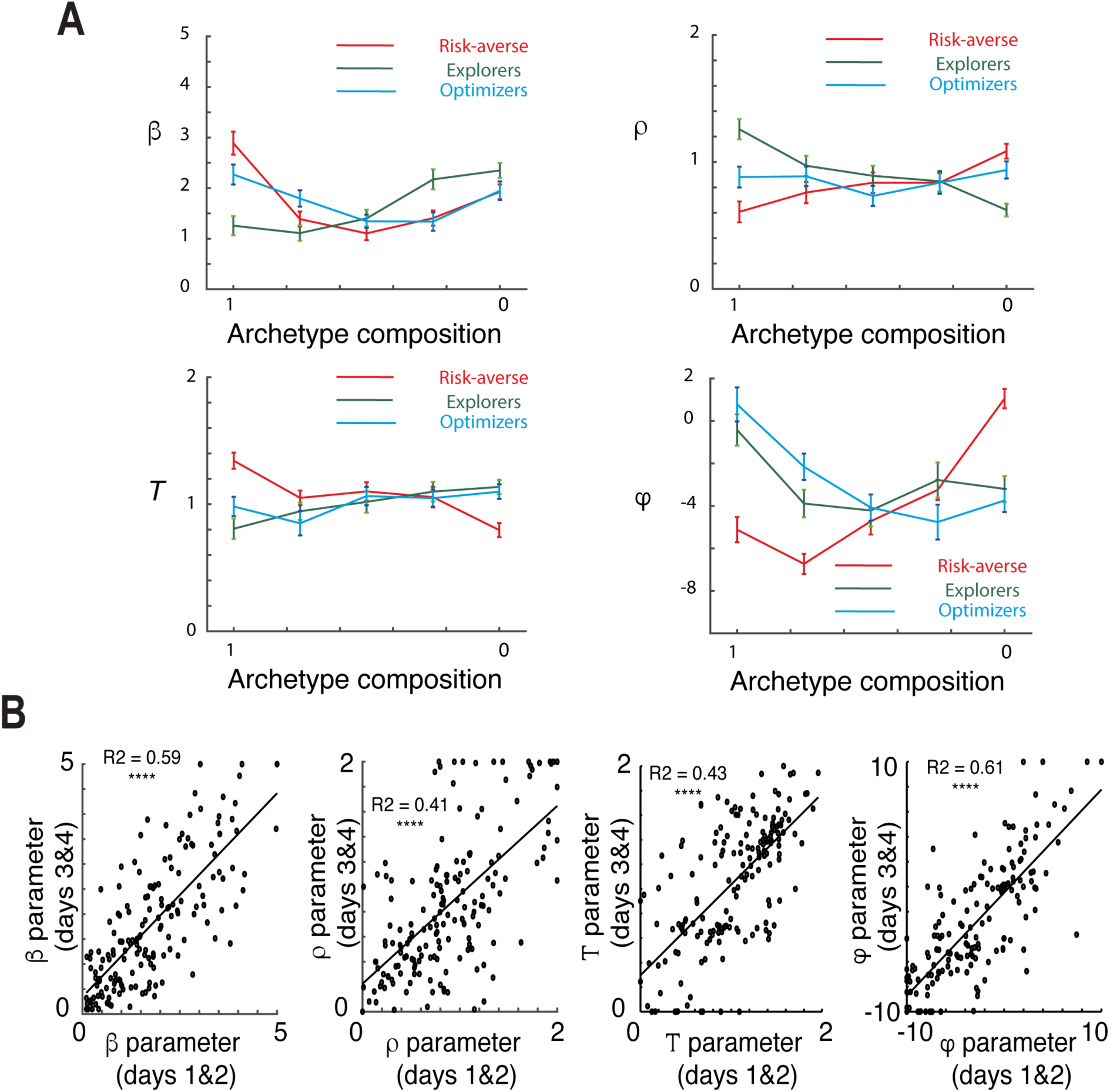
Additional model results. **A**. Model parameters as a function of archetypal compositions. **B**. Parameter stability was determined by comparing the parameters fitted over the first 2 days of baseline compared to parameters fitted over the last two days.

**Supplementary Figure 4.**
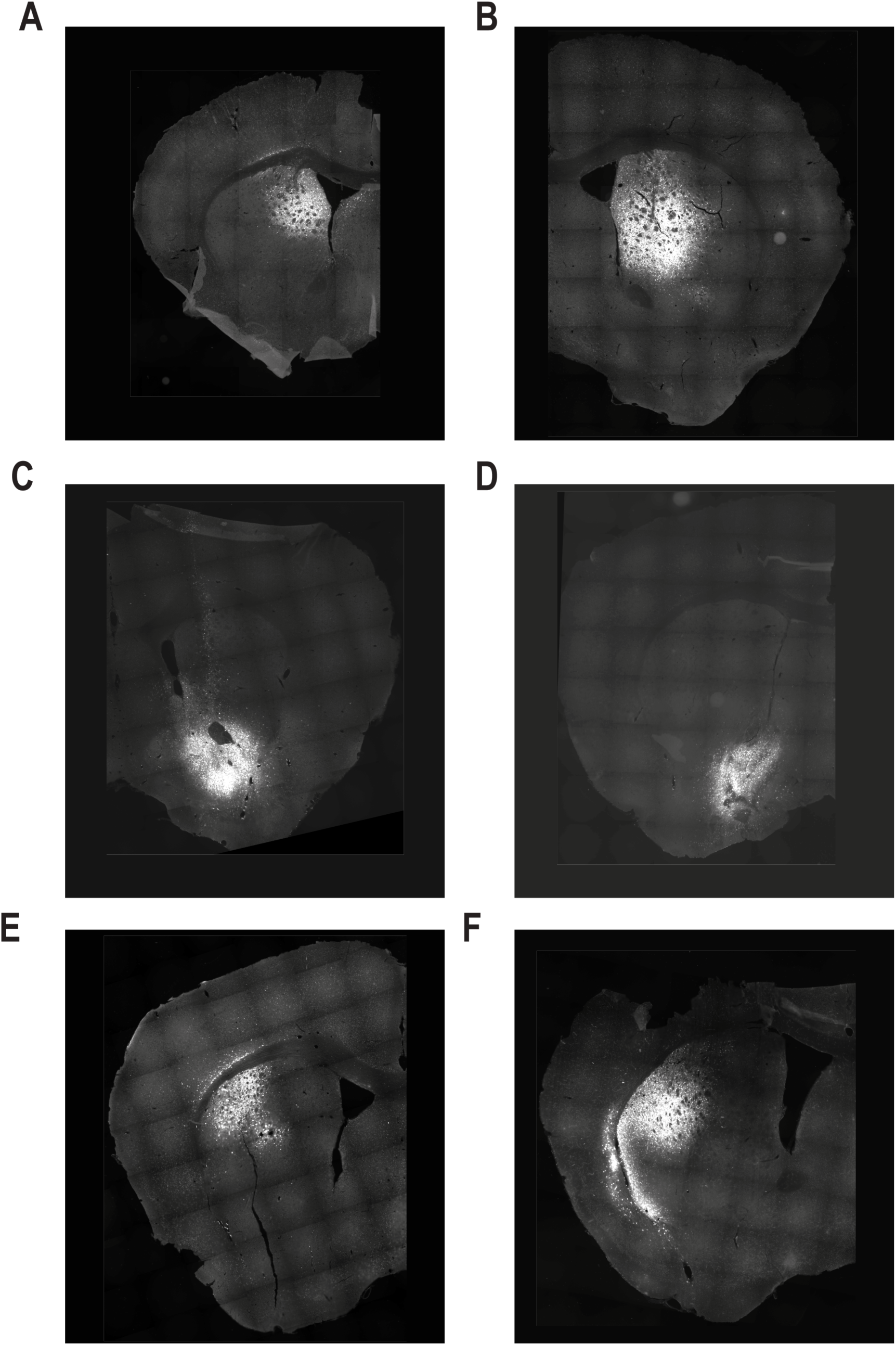
Initial targeting of three striatal regions. (A-B DMS, C-D NAc, and E-F DLS) in two mouse lines: Drd1-cre (A-C-E) and Adora2a-Cre (B-D-F), using the control viruses AAV5-hSyn1-DIO-mCherry.

**Supplementary Figure 5.**
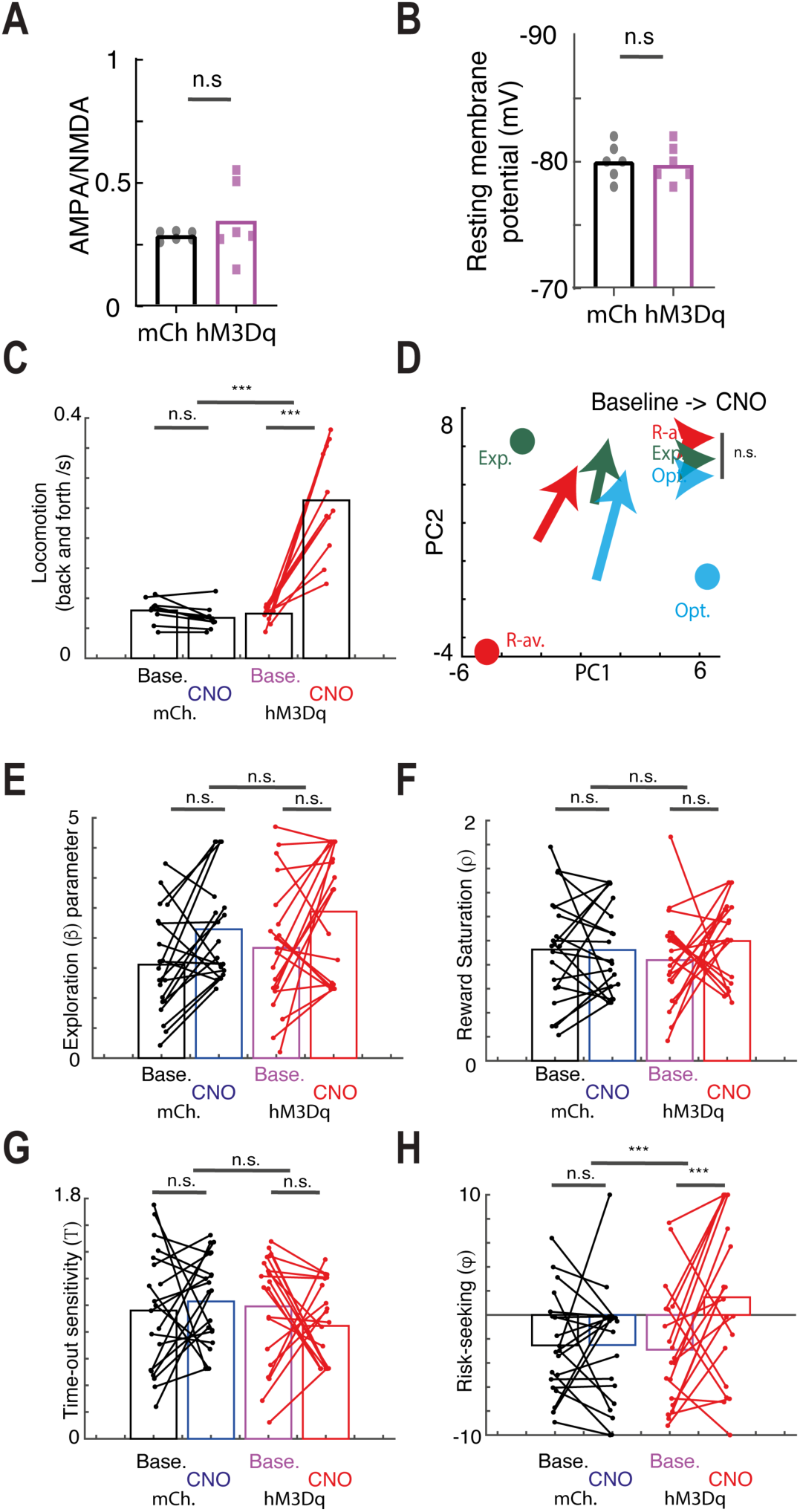
Additional measures and model for DMS-dSPN experiments. **A**. The AMPA/NMDA ratio was not different between conditions. **B**. The resting membrane potential was not different between conditions. **C**. Locomotion (back and forth movements in the conditioning box) for (baseline, CNO) x (mCherry, hM3Dq) conditions show a specific increase in locomotion following CNO on hM3Dq animals. **D**. Ternary plot showing the global effect of CNO on strategies split by archetypes, with each arrow showing group averages for each archetype from h3MDq animals under CNO. **E-H**. Model parameters (explore/exploit; reward saturation; time-out sensitivity; risk-seeking) for DREADD (hM3Dq) animals and mCherry (mCh) controls, under baseline and CNO conditions. Only the risk-seeking was significantly different in the (hM3Dq, CNO) condition.

**Supplementary Figure 6.**
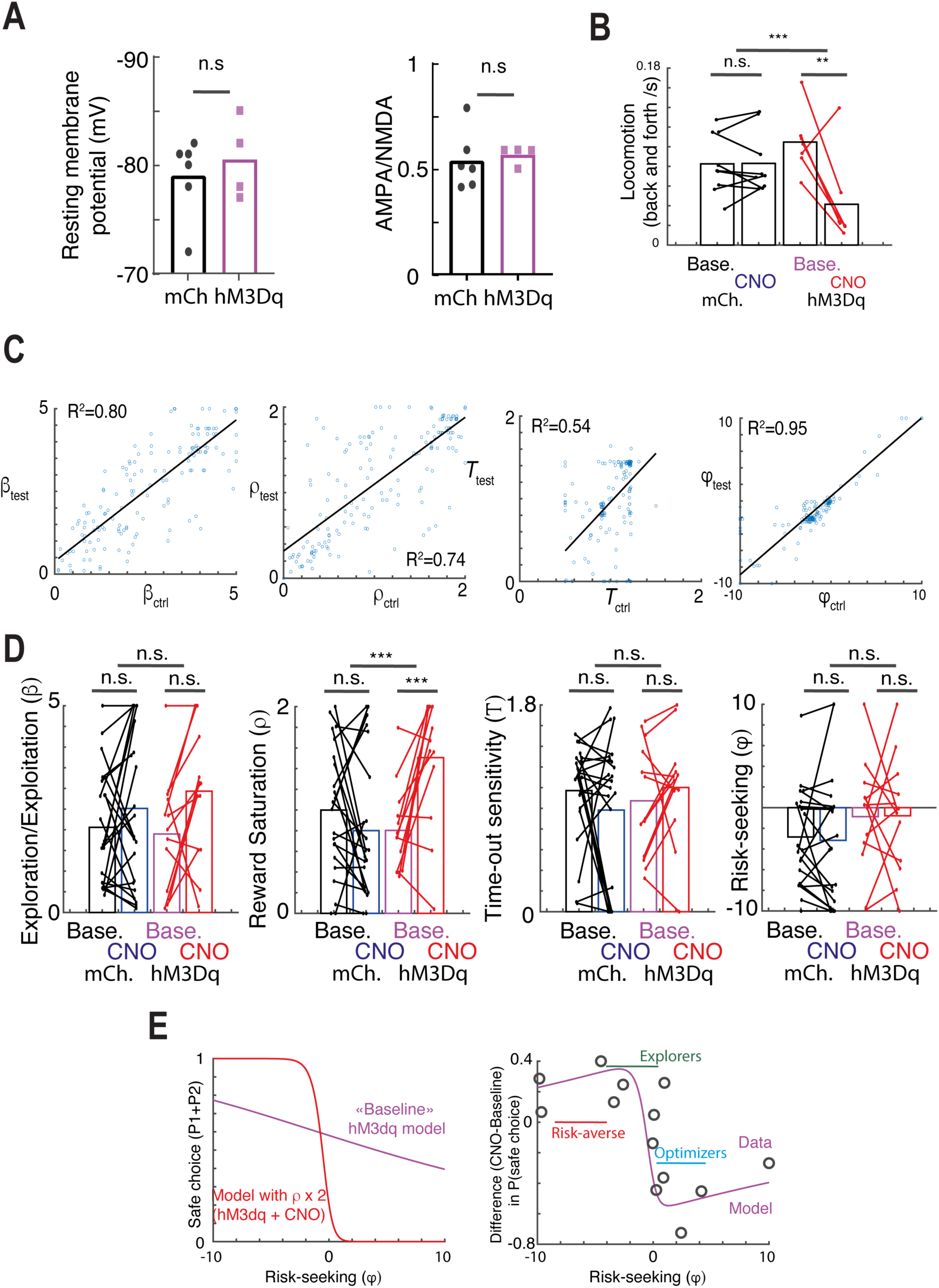
Additional measures and model for DMS-iSPN experiments. **A**. The the resting membrane potential (left) and the AMPA/NMDA ratio were not different between conditions. **B**. Locomotion (back and forth movements in the conditioning box) for (baseline, CNO) x (mCherry, hM3Dq) conditions show a specific decrease in locomotion following CNO on hM3Dq animals. **C**. Simulations showing the effects of a decrease in the number of trials on parameter recovery. Each dot depicts the parameter recovered from a simulation under the control number of trials (n=50) against the parameter recovered from a simulation under the h3Mdq number of trials. **D.** Model parameters (explore/exploit; reward saturation; time-out sensitivity; risk-seeking) for DREADD (hM3Dq) animals and mCherry (mCh) controls, under baseline and CNO conditions. Only the reward saturation was significantly different in the (hM3Dq, CNO) condition. **E**. Simulations of safe choices (P1+P2, as measured in Fig. 5) at baseline (purple) and following a decrease in reward saturation (increase in rho, red) depending on the risk seeking parameter (left); and data of safe choices under CNO for animals in the DMS-iSPN experiment against their risk-sensitivity parameter at baseline, showing a match with the model predictions (purple line, left).

**Supplementary Figure 7.**
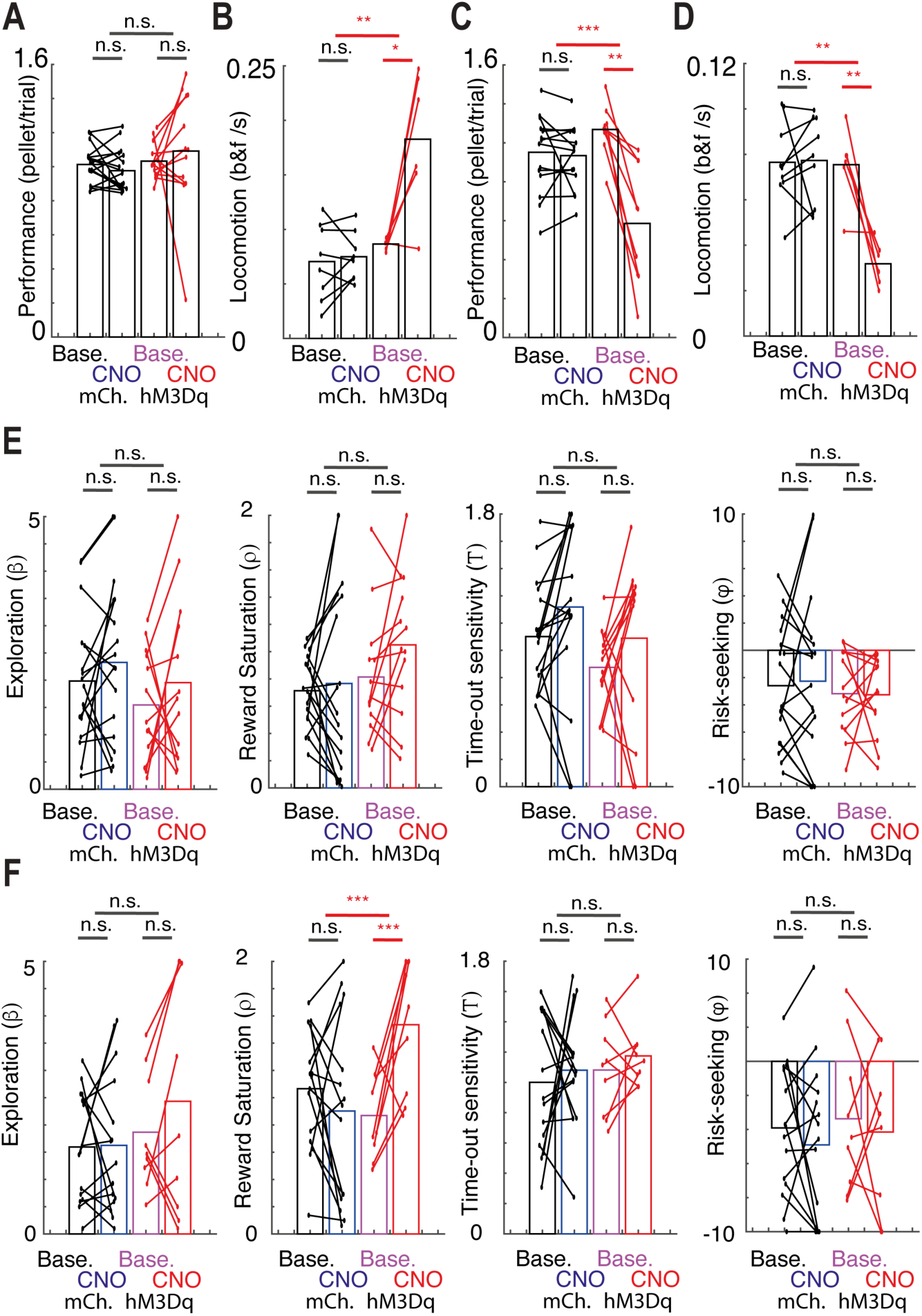
Additional measures and model for NAc experiments. **A.** CNO treatment decreased the performance in D1-NAc animals. **B.** Locomotion (back and forth movements in the conditioning box) for (baseline, CNO) x (mCherry, hM3Dq) conditions show a specific decrease in locomotion following CNO on A2A-NAc-hM3Dq animals. **C.** CNO treatment did not affect the performance in A2A-NAc animals **D.** Locomotion (back and forth movements in the conditioning box) for (baseline, CNO) x (mCherry, hM3Dq) conditions show a specific increase in locomotion following CNO on D1-NAc-hM3Dq animals. **E**. Model parameters (explore/exploit; reward saturation; time-out sensitivity; risk-seeking) for D1-NAc - DREADD (hM3Dq) animals and mCherry (mCh) controls, under baseline and CNO conditions. Only the risk-seeking was significantly different in the (hM3Dq, CNO) condition. **F**. Model parameters (explore/exploit; reward saturation; time-out sensitivity; risk-seeking) for A2A-NAc - DREADD (hM3Dq) animals and mCherry (mCh) controls, under baseline and CNO conditions, with no significant effect of CNO.

**Supplementary Figure 8.**
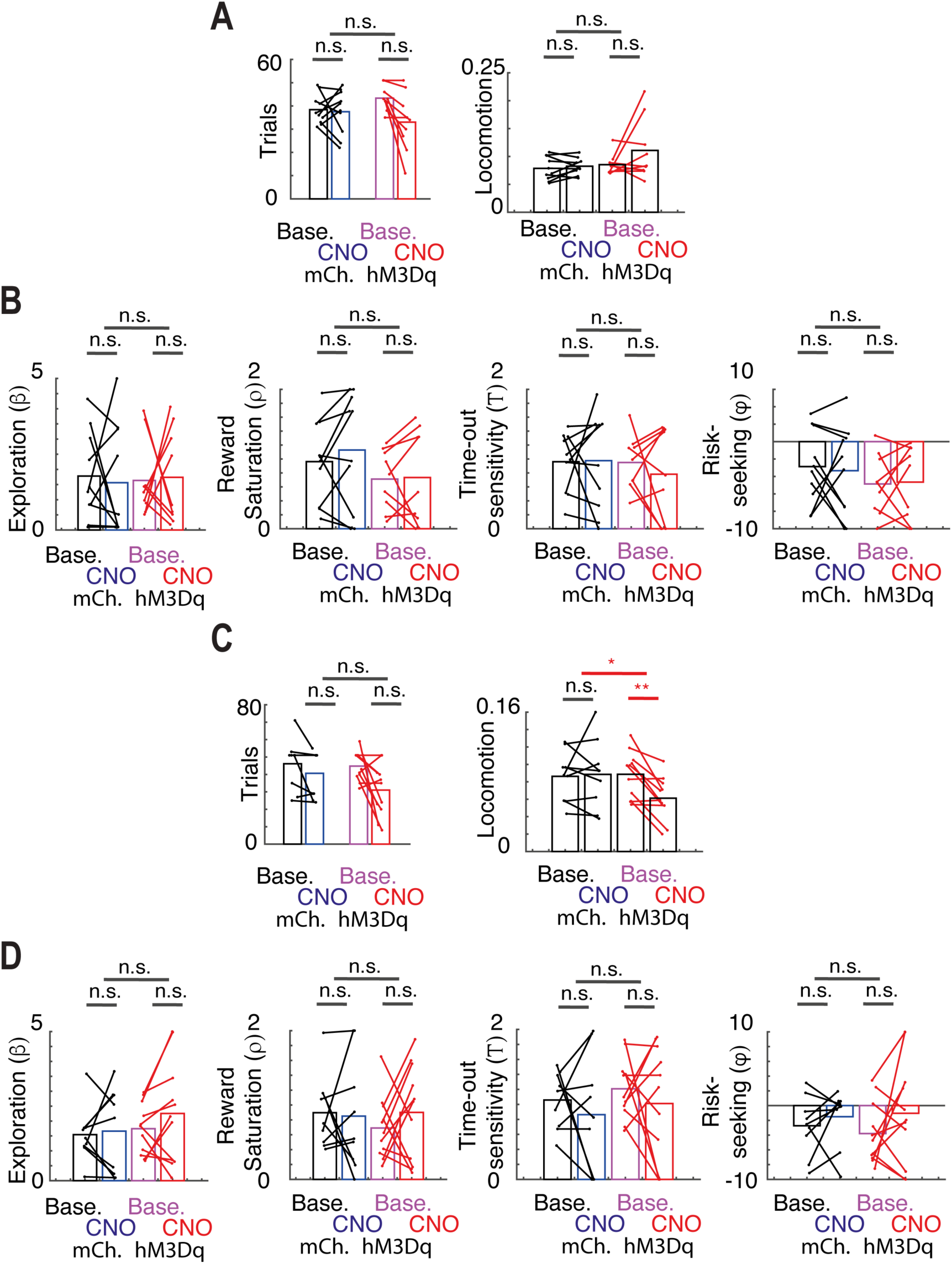
Model fits and behavioral measures for DLS animals. **A** model parameters (from left to right : explore/exploit ; reward saturation; time-out sensitivity; risk-seeking) and **B** behavioral measures (from left to right : number of trials, % of omissions, % of premature responses, and locomotion) for A2A-DLS (hM3Dq) animals and mCherry (mCh) controls, under baseline and CNO conditions. CNO only affected % of omissions (F_(1,18)_=5.32, p = 0.03; T_(10)_ = −5.63, p = 2.10^−4^), % of premature responses (F_(1,18)_=6.89, p = 0.02; T_(10)_ = 2.76, p = 0.02) and locomotion (F_(1,18)_=6.63, p = 0.02; T_(10)_ = −4.21, p = 0.002) in A2A-DLS-hM3Dq animals. (**C,D**) same for D1-DLS animals. CNO only affected % of omissions (F_(1,17)_=12.73, p = 0.02; T_(8)_ = −3.94, p = 0.04).

**Table 1.** Quantification of % of cFos-positive cells among mCherry positive cells.

|  | A2A |  | D1 |  |
| --- | --- | --- | --- | --- |
|  | hm3Dq | mCherry controls | hm3Dq | mCherry controls |
| DMS | 31.1% (SD=16.4, n=6) | 1.1% (SD=0.9, n=6) | 64.4% (SD=17.8, n=9) | 13.3% (SD=7.2, n=9) |
| DLS | 35.7% (SD=23.1, n=6) | 3.6% (SD=4.8, n=6) | 43.9% (SD=9.2, n=8) | 2.5% (SD=5, n=8) |
| NAC | 48.2% (SD=17.2, n=7) | 7.3% (SD=8.4, n=6) | 66.1% (SD=16.9%, n=7) | 13.9% (SD=15.9, n=7) |

